# A lettuce receptor-like kinase recognizes the highly conserved heptapeptide motif within microbial NEP1-like proteins

**DOI:** 10.1101/2025.11.10.687460

**Authors:** Iñigo Bañales, Sarah L. Mehrem, Samara Almeida Landman, Marrit Alderkamp, L.C.P. (Max) Pijfers, Stan Baijens, Alix von Bredow, Peter Schutte, Sander Prevoo, Gijs van Asselt, Sagayamary Sagayaradj, Basten Snoek, Richard Michelmore, Dmitry Lapin, Guido Van den Ackerveken

## Abstract

Plants rely on cell surface immune receptors to detect microbial patterns and initiate effective defense responses. Although the Asteraceae family is one of the largest and economically important plant groups, little information is available about its pattern-triggered immunity signaling. Cultivated lettuce (*Lactuca sativa* L.) recognizes a 24-amino acid peptide (nlp24) from necrosis- and ethylene-inducing peptide 1-like proteins (NLPs) found in bacteria, fungi, and oomycetes. Here, we perform an extensive characterization of nlp24-induced immune responses in lettuce and identify the LETTUCE nlp24 RECEPTOR (LNR) as the leucine-rich repeat receptor-like kinase mediating its recognition. Remarkably, nlp24 recognition in lettuce and subsequent activation of defenses strongly depend on the conserved heptapeptide motif (GHRHDWE). Structural modeling-guided mutagenesis experiments suggest that residues in the nlp24 heptapeptide interact with a hydrophobic pocket in the LNR solenoid structure. Divergent ligand specificities and the absence of sequence homology between LNR and *Arabidopsis* nlp24-recognizing receptor indicate that the NLP recognition in lettuce and *Arabidopsis* emerged independently, through convergent evolution. Our phylogenetic analysis shows that LNR is closely related to *Arabidopsis* MIK2 (MALE DISCOVERER 1-INTERACTING RECEPTOR-LIKE KINASE 2), but belongs to a distinct, Asteraceae-specific monophyletic subgroup that has undergone a significant expansion in *Lactuca*. Our findings provide insights into the mechanisms of pattern-triggered immunity in lettuce and the fast evolution of its immune receptor repertoire. On the translational side, our findings open opportunities for the crop defense improvement via interfamily transfer.

**Significance statement:** Understanding pathogen recognition in crops is key to improving disease resistance. Our study identified the cell surface immune receptor in cultivated lettuce that senses the nlp24 pattern derived from secreted proteins (NLPs) of prokaryotic and eukaryotic microbial pathogens. Unlike the previously characterized receptor from Arabidopsis (RLP23), the lettuce nlp24 receptor (LNR) detects the deeply conserved heptapeptide motif of NLP proteins that is required for host cell lysis. Transfer of the LNR receptor to a Solanaceous species conferred quantitative resistance to the oomycete pathogen *Phytophthora capsici*. Our findings advance the understanding of pattern-triggered immunity in a major leafy crop and highlight LNR as a promising receptor for broad-spectrum plant resistance engineering.

## Introduction

Immunity in plants is initiated by the recognition of pathogens by receptors located at the cell surface or inside of the cell (1–3). Receptors at the plasma membrane often perceive conserved molecules derived from or associated with microbial pathogen invasion (molecular patterns) (1, 2). These pattern-recognition receptors (PRRs) are divided into several classes, with the largest being leucine-rich repeat (LRR)-containing receptor-like kinases (RLKs) and receptor-like proteins (RLPs), which possess or lack a cytoplasmic kinase domain, respectively (1, 2). LRR-RLPs constitutively interact with the adapter RLK SOBIR1 (SUPPRESSOR OF BRASSINOSTEROID INSENSITIVE 1 (BRI1)-ASSOCIATED KINASE (BAK1)-INTERACTING RECEPTOR KINASE 1) to form functional signaling complexes (4, 5). Upon ligand binding, LRR-RLPs and LRR-RLKs typically associate with the co-receptor BAK1 (BRASSINOSTEROID-INSENSITIVE 1 (BRI1) ASSOCIATED RECEPTOR KINASE 1) or other members of the SERK (SOMATIC EMBRYOGENESIS RECEPTOR KINASE) family (4, 5). Functional outputs of pattern-triggered immunity (PTI) include calcium influx and reactive oxygen species (ROS) burst, activation of mitogen-activated protein kinase (MPK) via phosphorylation, and host transcriptional reprogramming (6, 7). PRRs are also required for robust plant immunity conferred by intracellular receptors (8–10).

A few PRRs are relatively conserved in the plant kingdom. These conserved PRRs are exemplified by FLS2 (FLAGELLIN-INSENSITIVE 2) and CERK1 (CHITIN ELICITOR RECEPTOR KINASE 1), which recognize the bacterial flagellin epitope flg22 (11, 12) and fungal chitin (13–15), respectively. Similarly, the immunity- and growth-related MIK2 (MALE DISCOVERER 1-INTERACTING RECEPTOR-LIKE KINASE 2) receptor in *Arabidopsis thaliana* (*Arabidopsis* hereafter) has orthologs outside Brassicaceae (16). However, often PRRs belong to expanded, fast-evolving groups of RLPs and RLKs, and they are restricted to specific taxonomic groups such as families or genera (1), for instance EFR (ELONGATION FACTOR-TU RECEPTOR) (17) and RLP32 (18) in Brassicaceae and the Pep-13 receptor unit (PERU) and the EIX receptor 2 (EIX2) in Solanaceae (19, 20). Despite between-species variation in PRR repertoires, convergent evolution for the recognition of molecular patterns is also observed. For instance, RLP30 and RE02 are sequence-unrelated RLPs sensing small cysteine-rich proteins (SCPs) in *Arabidopsis* and *Nicotiana benthamiana*, respectively (21). Similarly, elicitins are recognized by evolutionarily distant RLPs in Solanaceae species *N. benthamiana* and *Solanum microdontum* (22).

Patterns triggering PRR-mediated immunity are usually restricted to individual microbial groups such as bacteria (23–25). However, necrosis- and ethylene-inducing peptide 1 (Nep1-like proteins or NLPs) are secreted by bacteria, fungi, and oomycetes (26–29). NLPs are found in major groups of plant pathogens with different lifestyles, including hemibiotrophic *Fusarium* and *Phytophthora* and biotrophic downy mildew species (26, 30). A subset of NLPs functions as cytolytic toxins, causing host cell collapse within minutes after exposure (31). The structure of NLPs resembles that of actinoporins, and, similarly, they can form oligomeric complexes bound to the membranes, resulting in the lysis of plant cells (32–34). A conserved heptapeptide motif in the center of the protein is crucial for the cytotoxic activity of NLPs (GHRHDWE) (34, 35). Residues within this motif are part of the cation-binding pocket that is essential for the interaction with membrane sphingolipids and subsequent cell lysis (34, 35). Interestingly, the heptapeptide motif is conserved across all phylogeny-based NLP groups (types I-III), regardless of their cytotoxicity (26, 36), suggesting functions beyond the cytotoxic activities. Damage caused by the cytolytic activity of NLPs leads to the activation of plant immune responses (34, 37). However, NLPs can also activate defense outputs independently of their cell-death triggering activity (36, 38).

A short peptide of 24 amino acids (nlp24) derived from *Arabidopsis* downy mildew NLP3 protein or nlp20 from *Phytophthora parasitica* are strong elicitors of immunity in Brassicaceae species (28, 39, 40). The nlp24 sequence contains two major conserved regions: the first N-terminal 11 residues, conserved in type I NLPs (region I) (28, 36), and the C-terminal heptapeptide (region II) (36, 38). In nlp20, the heptapeptide motif is shortened (38). In *Arabidopsis*, nlp24 is recognized by the cell surface receptor RLP23, which shows specificity towards the AIMY residues within region I (41). Nlp24 is also recognized in distantly related species, such as cultivated lettuce (*Lactuca sativa* L.) from Asteraceae (38). Since the RLP23 clade is absent from the lettuce reference genome (42), lettuce is expected to perceive nlp24 via a different receptor.

Here, we identified the receptor responsible for nlp24 recognition in lettuce and characterized the nlp24-triggered PTI in this species. Nlp24 treatment induced the production of ROS species, phosphorylation of MPK3 and 6 (MITOGEN-ACTIVATED PROTEIN KINASES), and immunity-related transcriptional reprogramming. Removal of the heptapeptide motif resulted in a loss of immunity-eliciting activity. Screening for nlp24-triggered ROS burst in a *L. sativa* diversity panel, Genome-wide Association Studies (GWAS), and comparison of whole-genome assemblies of responsive and non-responsive cultivars uncovered a MIK2-like LRR-RLK as a receptor candidate. Transient complementation assays in lettuce and *N. benthamiana* confirmed it as the *LETTUCE nlp24 RECEPTOR* (*LNR*), the first cell surface immune receptor reported in lettuce and in any member of the Asteraceae family of plants. LNR interfamily transfer to Solanaceae conferred increased resistance to a *Phytophthora* pathogen. Structural modeling-guided mutagenesis supported a direct interaction between LNR and residues in the nlp24 heptapeptide. Phylogenetic analysis showed that LNR belongs to an Asteraceae-specific monophyletic clade that has undergone a significant expansion in *Lactuca* and forms a sister group to orthologs of *Arabidopsis* MIK2. Our study provides insights into the rapid evolution of the cell surface immune receptors in lettuce, identifies a receptor of a broadly present microbial molecular pattern, and provides a basis for future work on PTI mechanisms in Asteraceae and translational research for crop engineering.

## Results

### Nlp24 activates lettuce immunity through a different receptor than in *Arabidopsis*

The nlp24 peptide (Figure 1a, representative nlp sequence from *Hyaloperonospora arabidopsidis* NLP3) acts as an immunogenic pattern in lettuce (*Lactuca sativa* L.), as it provided acquired resistance against lettuce downy mildew *Bremia lactucae* (38). To gain a better understanding of the underlying molecular events, we characterized early nlp24-triggered immunity outputs in lettuce. The nlp24 treatment resulted in a ROS burst in five out of nine tested lettuce cultivars, including cv. Olof, with a peak at ∼7 minutes (Figure 1b, Supplementary Figure 1). Similarly, phosphorylation of the MAP kinases MPK3 and MPK6 was detectable at 40 minutes (Figure 1c). Transcriptome profiling of nlp24-treated leaf tissue of cv. Olof (1h after infiltration) revealed 154 upregulated and 14 downregulated genes (Figure 1d, Supplementary File 1, |log2FC|=>1, adjusted p-value<=0.05). The nlp24-upregulated lettuce genes were enriched for GO terms related to plant immunity, such as ‘innate immune response’, ‘defense response to symbiont’, or ‘plant-type hypersensitive response’ (Supplementary Figure 2, Supplementary File 2). Similarly, there was a significant overlap between the upregulated lettuce genes and *Arabidopsis* genes induced during PTI (Fisher’s exact test, p-value=2.2 * 10^-16^, odds ratio = 11.9) (43). Furthermore, WRKY transcription factor (TF) DNA binding motifs were significantly enriched in the promoters of the upregulated genes (Supplementary File 3) (44, 45). Overall, these features demonstrate that nlp24 acts as an immunogenic microbial pattern in lettuce.

**Figure 1.**
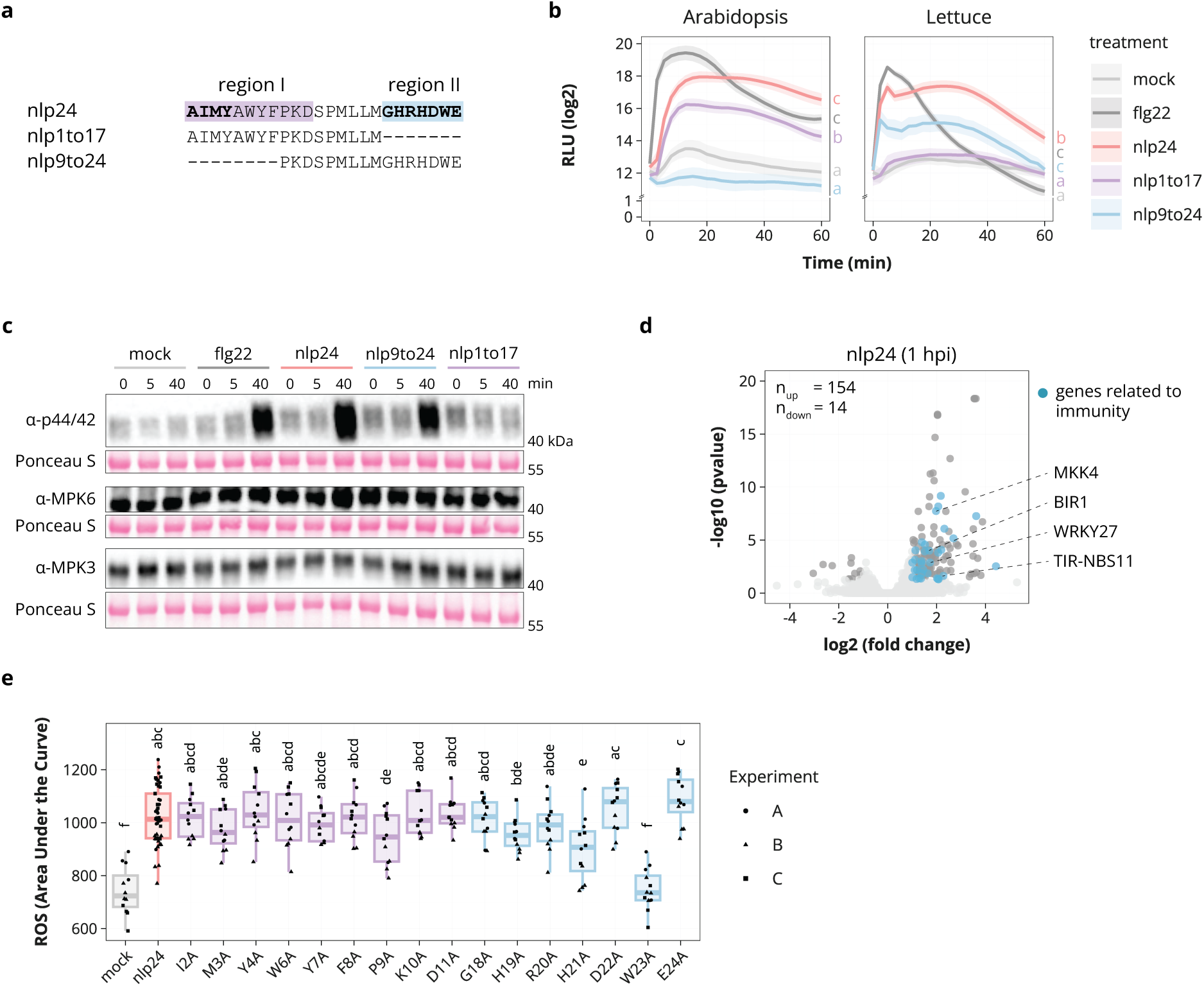
Nlp24 peptide acts as an immunogenic pattern in lettuce. **a.** Amino acid sequence alignment of nlp24, nlp1to17, and nlp9to24. Conserved regions I (purple) and II (blue) are highlighted at the N- and C-termini, respectively. **b.** Apoplastic reactive oxygen species (ROS) production in *Arabidopsis* Col-0 and lettuce cv. Olof in response to flg22 (0.2 µM), nlp24 (2 µM), nlp1to17 (2 µM), nlp9to24 (2 µM), or mock (0.02 % DMSO). The ROS burst was measured over 1 hour. Mean and standard error per time point is shown (n=8, two independent experiments). Letters represent levels of statistical significance between mean area under the curve (AUC) values (ANOVA, Tukey’s honestly significant difference (HSD), p<0.05). **c.** MPK activation in lettuce cv. Olof treated with the same elicitors as in **b**. Samples were harvested immediately (0), 5, and 40 minutes after treatment. Phosphorylated MPKs were detected by immunoblots with α-p44/42-ERK antibody. MPK3 and MPK6 levels in the same samples were detected with α-MPK3/6 antibodies. Ponceau S staining of the large RUBISCO subunit serves as a loading control. The experiment was repeated two times with similar results. **d.** Volcano plot of differentially expressed genes (|log2FC|=>1, adjusted p-value<=0.05) in lettuce cv. Olof 1 hour after treatment with nlp24 (0.5 µM) compared to mock (MQ). Genes with a hit to *Arabidopsis* sequences annotated with immunity-related GO terms are colored in blue. Selected immunity-related genes are labeled. **e.** ROS burst in lettuce cv. Olof treated with nlp24 (2 µM), alanine substitution variants (2 µM), or mock (0.02 % DMSO). ROS curves over 73.5 minutes were translated to AUC values. Letters represent levels of statistical significance (ANOVA, Tukey’s HSD, p<0.05; n=48 for nlp24, n=12 for alanine substitution variants; three independent experiments).

In *Arabidopsis*, the cell surface-localized receptor-like protein RLP23 recognizes nlp peptides via the AIMY motif within the N-terminal region I of nlp24 (Figure 1a, (41)); however, this specificity was not established for lettuce. Remarkably, the nlp24 variant without this region (nlp9to24, Figure 1a) induced a ROS burst in lettuce cv. Olof, yet with a lower intensity than the full-length nlp24 (Figure 1b, Supplementary Figure 1). As expected, the nlp9to24 peptide treatment did not produce a ROS burst in *Arabidopsis* (Figure 1b, Supplementary Figure 1). The nlp9to24 peptide also triggered MPK3 and MPK6 phosphorylation in lettuce (Figure 1c). An nlp24 variant lacking the conserved heptapeptide motif in the C-terminal region II (nlp1to17, Figure 1a) was not immunogenic in lettuce at the level of the ROS burst and MPK3 and MPK6 phosphorylation, whereas it did trigger the ROS burst in *Arabidopsis* consistent with earlier reports (Figure 1b, c, Supplementary Figure 1). An alanine substitution screen confirmed that the heptapeptide motif (GHRHDWE) is essential for nlp24 recognition in lettuce, since the substitutions H21A and W23A partially or completely abolished the ROS burst, respectively (Figure 1e, Supplementary Figure 3). To conclude, differences in the recognition specificities between *Arabidopsis* and lettuce as well as the previously reported absence of *RLP23* homologs in the lettuce reference genome (42) suggest convergent evolution of the NLP recognition capacity in these two species.

### NLP recognition in lettuce is largely driven by its conserved heptapeptide motif

To test whether NLPs from lettuce-adapted pathogens can be recognized by lettuce, we performed ROS burst assays with nlp24-like peptides from the oomycete *Bremia lactucae*, the fungus *Fusarium oxysporum* f. sp. *lactucae*, and the bacteria *Pectobacterium carotovorum* and *Bacillus subtilis*. Ten out of 16 tested peptides produced nlp24-like ROS burst curves (Figure 2a, b, Supplementary Figure 4). The *P. parasitica* nlp20 did not induce a ROS burst in lettuce (Figure 2a, b, Supplementary Figure 4). Variation in the heptapeptide motif in these sequences could not be readily linked with differences in the ROS bursts, suggesting that amino acids outside the heptapeptide motif contribute to nlp24 recognition in lettuce, even though the heptapeptide is the main determinant of the recognition (Figure 1b, c, e, Supplementary Figure 3). To gain a better understanding of NLP recognition via the heptapeptide motif and other sequences in lettuce, we analyzed the ROS burst in response to nlp24 variants with different lengths of the heptapeptide (Figure 2c, d, Supplementary Figure 5). A ROS burst was triggered with full-length nlp24 and heptapeptide-only fragment (nlp18to24) but not with variants lacking the heptapeptide motif (nlp1to11, nlp1to13, nlp1to17) or its last three amino acids (DWE, nlp1to21) (Figure 2c, d, Supplementary Figure 5). At the same time, variants with only the first two amino acids of the heptapeptide (nlp1to19) or missing the last two (nlp1to22) were still recognized (Figure 2c, d, Supplementary Figure 5). Interestingly, simultaneous deletion of the AIMY motif and the last two amino acids in the heptapeptide motif (nlp9to22 and nlp5to22) resulted in the loss of the ROS burst response (Figure 2c, d, Supplementary Figure 5). These results demonstrate that the efficient NLP recognition in lettuce depends mainly on its conserved heptapeptide motif, but other molecular features contribute as well, particularly when the heptapeptide motif is mutated.

**Figure 2.**
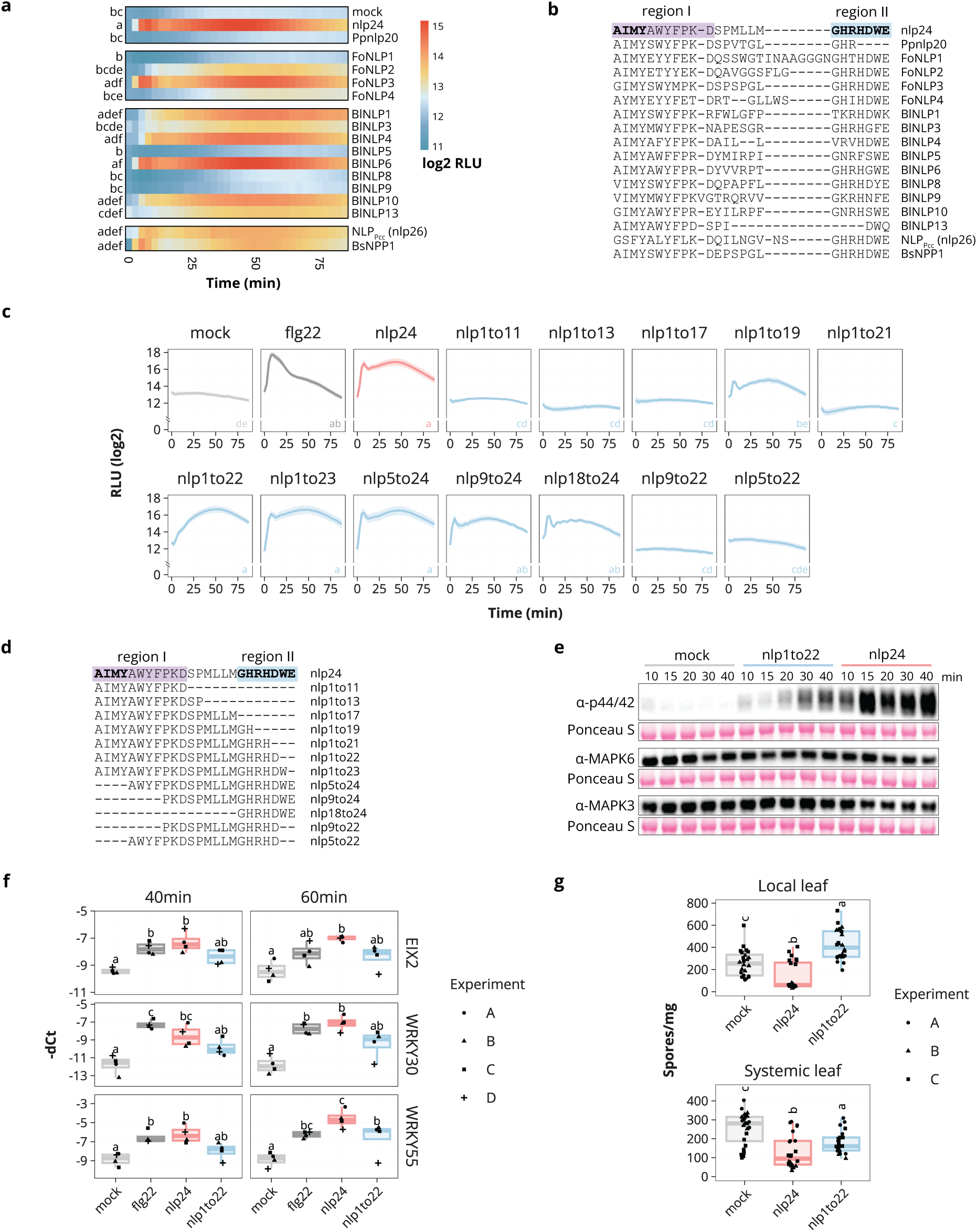
Conserved heptapeptide motif in nlp24 can activate robust immunity signaling in lettuce. **a.** ROS burst in lettuce cv. Olof in response to nlp24 (2 µM), or nlp24-like peptides from *Fusarium oxysporum* f. sp. *lactucae* (Fo), *Bremia lactucae* (Bl), *Pectobacterium carotovorum* (Pcc) and *Bacillus subtilis* (Bs) (2 µM), nlp20 from *Phytophthora parasitica* (Pp), or with mock (0.02 % DMSO). The ROS burst was measured over 82.5 minutes. Mean values per time point are shown (n=15 for mock, n=16 for nlp24, n=6-8 for nlp24-like peptides; two independent experiments). Letters represent levels of statistical significance between mean AUC values (ANOVA, Tukey’s HSD, p<0.05). **b.** Alignment of nlp24 and nlp24-like peptide sequences from pathogens. **c.** ROS burst in lettuce cv. Olof in response to nlp24 and variants of different length (2 µM) measured over 87.5 minutes. Mean and standard error per time point is shown (n=8; two independent experiments). Letters represent levels of statistical significance between mean AUC values (ANOVA, Tukey’s HSD, p<0.05). **d.** Alignment of nlp24 and truncated variants. **e.** MPK activation in lettuce cv. Olof treated with mock (0.02 % DMSO), nlp24 (2 µM), or nlp1to22 (2 µM) over multiple time points (10-40 minutes). Phosphorylated MPKs were detected by immunoblots with α-p44/42-ERK antibody. MPK3 and MPK6 levels in the same samples were detected with α-MPK3/6 antibodies. Ponceau S staining of blots serves as a loading control. The experiment was repeated two times with similar results. **f.** Expression levels of *EIX2*, *WRKY30*, and *WRKY55* in lettuce cv. Olof after treatment with mock (0.02 % DMSO), flg22 (0.2 µM), nlp24 (2 µM), or nlp1to22 (2 µM). Letters represent levels of statistical significance (ANOVA, Tukey’s HSD, p<0.05; n=4). **g.** Sporulation levels of *Bremia lactucae* Bl:21 isolate on lettuce cv. Olof. The first true leaves were infiltrated with elicitor (2 µM; 0.02 % DMSO as mock) and inoculated with the pathogen 24 hours later. Spores were quantified at 13 dpi and normalized to leaf weight. Letters indicate levels of statistical significance (ANOVA, Tukey’s HSD, p<0.05; n=48; three independent experiments).

To investigate further the importance of the intactness of the heptapeptide for the immune system activation in lettuce, we compared immunity outputs after exposing lettuce to the nlp24 full-length and nlp1to22 variants. The C-terminally truncated variant did not give the early ROS burst peak (at ∼7 min) observed with the full-length nlp24 (Figure 2c, d, Supplementary Figure 5, nlp1to22). Consistently, MPK3 and MPK6 phosphorylation was delayed by ∼10 minutes and was weaker overall (Figure 2e). Analysis of nlp24 marker gene expression and local and systemic acquired resistance to *B. lactucae* induced by the peptide pretreatment confirmed that nlp1to22 is a weak elicitor of immunity (Figure 2f, g). Thus, the NLP recognition in lettuce is driven primarily by the conserved heptapeptide motif (GHRHDWE), which needs to be complete for the robust activation of immunity.

### Genome-wide association mapping of the nlp24-triggered ROS burst in lettuce

Of the nine lettuce cultivars initially tested in the ROS burst assays with nlp24, five responded to the peptide (Supplementary Figure 1), suggesting that one could use genetic and phenotypic variation in the lettuce germplasm to identify the nlp24 receptor gene. For this, we conducted a Genome-Wide Association Study (GWAS) using a diversity panel of 198 *L. sativa* accessions (46–48). An apoplastic ROS burst after exposure of leaf discs to nlp24 was detected in 101 of the 198 accessions (Figure 3a), but the genome reference cv. Salinas and the wild relative genome reference lines (*L. serriola* CGN25282, *L. saligna* CGN05327, *L. virosa* CGN04683) showed no ROS burst response (Supplementary Figure 6). Principal component analysis (PCA) of features extracted from the ROS curves confirmed a split between responsive and non-responsive accessions (Supplementary Figure 7), although quantitative differences in the ROS production between responsive accessions were detected too (Figure 3a, Supplementary Figure 7). The nlp24-responsive accessions came from Europe and Asia, without a clear geographical signature (Supplementary Figure 8). Segregation analysis of the nlp24 recognition at the ROS burst level in an F_2_ population from a cross between a responsive accession (cv. Crna) and the non-responsive reference line (cv. Salinas) suggested it is governed by a single dominant locus (χ^2^ = 1.333, p-value = 0.248) (Figure 3b).

**Figure 3.**
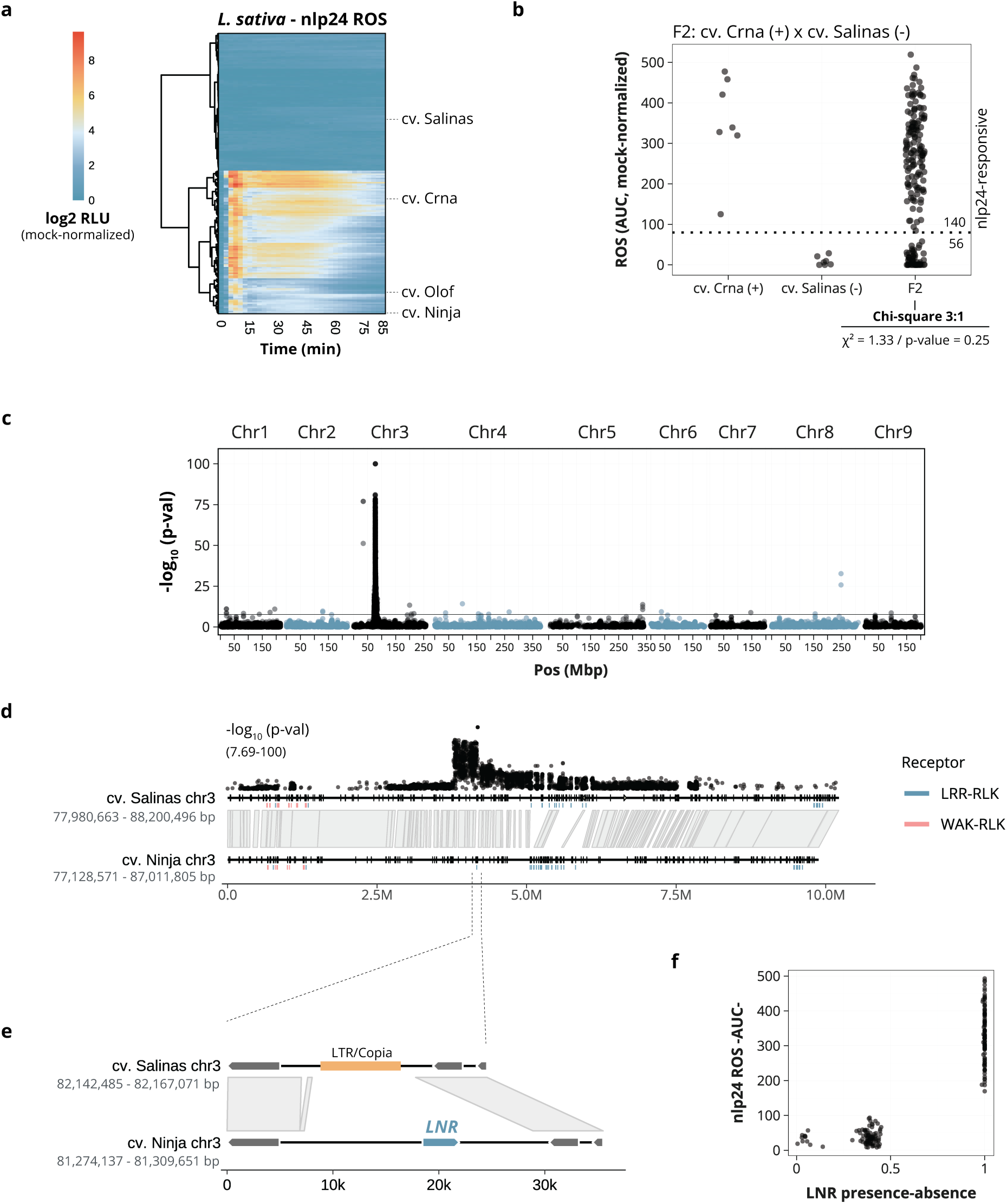
Mapping of the *LETTUCE nlp24 RECEPTOR* (*LNR*). **a.** ROS burst in 198 *L. sativa* accessions. Leaf discs were exposed to nlp24 (2 µM) or mock (0.02 % DMSO). Mean values over time are shown, normalized to mock (n=5-6, n=4 for CGN20716 and CGN14606). Accessions mentioned in the text are highlighted. **b.** ROS burst in F_2_ progeny (72.5 min) from a cross between cv. Crna (+) and cv. Salinas (-) in response to nlp24 (2 µM) (196 individuals). AUC values were normalized to mock-treated parental lines. F2 segregation fit a 3:1 ratio (χ2 = 1.33, p-value = 0.25). **c.** Manhattan plot after genome-wide association study (GWAS) for nlp24-triggered ROS burst (AUC). The x-axis shows genomic position across the *L. sativa* cv. Salinas genome (GCF_002870075.3, v8, Mb), with chromosome names above. The y-axis contains the −log_10_ p-values for the association of each SNP with the phenotype. The horizontal line shows the Bonferroni-adjusted significance threshold (-log_10_(P)>7.69). **d.** Comparison of the locus on chromosome 3 containing SNPs significantly associated with peptide recognition between cv. Salinas (GCF_002870075.4, v11) (-) and cv. Ninja (+). SNPs and their –log₁₀(p) values from GWAS are shown above. Positions of LRR-RLKs (blue) and WAK-RLKs (red) are indicated. Homology blocks (grey) were calculated with Minimap2. **e.** Zoom-in of the region containing the *LETTUCE nlp24 RECEPTOR* (*LNR*) gene. The LTR/Copia element in cv. Salinas is marked in orange. **f.** *LNR* presence-absence variation in 198 *L. sativa* accessions. The x-axis contains *LNR* coverage from whole-genome resequencing data (46, 47) aligned to the cv. Ninja genome, and the y-axis shows ROS score (AUC) after nlp24 treatment.

GWAS with a whole-genome single-nucleotide polymorphism (SNP) matrix derived from resequencing data (46–50) revealed a significant association of the nlp24-triggered ROS burst with chromosome 3 between positions 77,980,663 – 88,200,496 bp (Figure 3c, cv. Salinas genome v11 coordinates). This interval contains 258 genes in the lettuce reference genome (v11), including 32 receptor-like kinases (RLKs), which are organized into clusters (Figure 3d). SNP profiles across the lettuce panel suggest two haplotypes within this region, with non-reference haplotype correlating with the ROS burst (Supplementary Figure 9). A comparison of the locus sequence in the nlp24-non-responsive reference cv. Salinas and the responsive cv. Ninja confirmed the association between the nlp24 recognition and the non-reference haplotype in cv. Ninja (Figure 3d, e). The two haplotypes are mostly collinear except for the LRR-RLK cluster in the center of the locus (Figure 3d, Supplementary Figure 10). One LRR-RLK gene (*LSAT002_V1C300114880*) in the region with the strongest association to the nlp24-induced ROS burst was unique to cv. Ninja, suggesting that it encodes the putative receptor, which we tentatively called LETTUCE nlp24 RECEPTOR (Figure 3d, e, LNR). Analysis of the *LNR* presence-absence variation across the 198 lettuce accessions using short-read DNA sequences revealed that only the nlp24-responsive genotypes contain *LNR*, and that they all share the same sequence (Figure 3f), making *LNR* the prime candidate for the nlp24 receptor-encoding gene.

### *LNR* encodes an LRR-receptor-like kinase mediating NLP heptapeptide recognition in lettuce

LNR has the characteristic domain architecture of receptor-like kinases with an LRR-type ectodomain, a transmembrane region, and a cytoplasmic kinase domain with an RD-type catalytic site (Figure 4a). To test if LNR is responsible for the nlp24 recognition in lettuce, we performed transient complementation assays in leaves of the nlp24-non-responsive cv. Salinas. Ectopic expression of an *LNR* paralog from cv. Salinas (*LNR* paralog, negative control) did not result in the nlp24-induced ROS burst (Figure 4b, Supplementary Figure 11). Conversely, transient expression of *LNR* produced a ROS burst in response to full-length nlp24 but not its variants without the heptapeptide (nlp1to17) or with the W23A substitution (Figure 4b, Supplementary Figure 11). A similar pattern was observed when *LNR* and the cv. Salinas *LNR* paralog (negative control) were transiently expressed in *N. benthamiana* leaves (Supplementary Figure 12). Transient expression of *LNR* in *N. benthamiana* also resulted in reduced disease symptoms after the infection with *P. capsici* (LT3112 isolate), whose genome encodes 39 NLPs, many of them with the conserved heptapeptide motif (Figure 4c (51)). Substitution of the catalytic aspartate residue by asparagine in the kinase domain of LNR (Figure 4a, D958N) resulted in a significant loss of ROS burst activity in lettuce after nlp24 treatment, and the addition of C-terminal myc or GFP tags made this effect more pronounced (Figure 4g, Supplementary Figure 13). Taken together, these results show that LNR is the nlp24-recognizing RLK in lettuce requiring a functional cytoplasmic kinase domain for its full activity.

**Figure 4.**
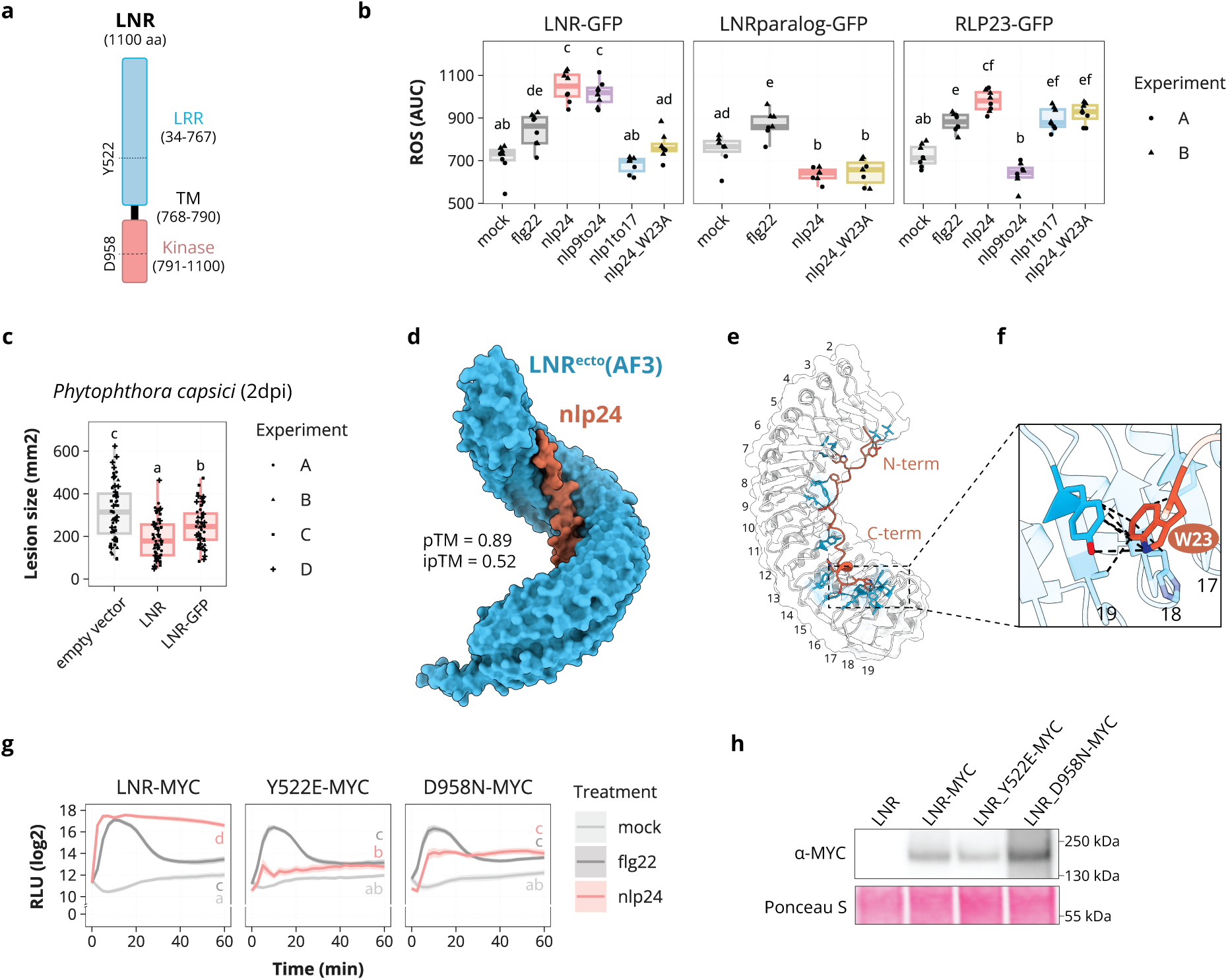
LNR is the RLK mediating NLP recognition in lettuce. **a.** Scheme of LNR protein domains, including the LRR-type ectodomain, the transmembrane domain, and the intracellular kinase domain. Amino acid coordinates are indicated in the brackets, and approximate locations of residues mentioned in the text are highlighted. **b.** ROS burst in lettuce cv. Salinas transiently expressing GFP-tagged versions of *RLP23*, *LNR*, and a closely related *RLK* (*LNR paralog*). Leaf discs were exposed to distinct elicitors 3 days after *Agrobacterium* infiltration. ROS burst was measured over 60 minutes. Letters represent levels of statistical significance between mean AUC values for all constructs (ANOVA, Tukey’s HSD, p<0.05; n=7-8; two independent experiments). **c.** *Phytophthora capsici* infection (LT3112 isolate) of *N. benthamiana* transiently expressing an empty vector, *LNR*, or *LNR-GFP*. Mycelium plugs were put on top of detached leaves 3 days post *Agrobacterium*-mediated transformation, and infection was assessed 2 days post infection. Lesion size (mm2) was measured from RGB pictures. Letters correspond to statistical significance levels among mean infection values (ANOVA, Tukey’s HSD, p<0.05; n=54-56; four independent experiments). **d.** Interaction between LNR ectodomain and nlp24 modeled by AlphaFold3. pTM and ipTM scores are included. **e.** Close-up view of the LRRs in contact with nlp24 as predicted by AlphaFold3 (LRRs 2-19). LNR residues predicted to interact with nlp24 are colored in blue, numbers on the convex side refer to the LRR numbering. **f.** Zoom-in of the predicted interaction between LNR^Y522^ and nlp24^W23^. **g.** ROS production in lettuce cv. Salinas transiently expressing MYC-tagged versions of *LNR*, *LNR^Y522E^*, or *LNR^D958N^*, exposed to various elicitors. Leaf discs were exposed to distinct elicitors 3 days after *Agrobacterium* infiltration. ROS burst was measured over 60 minutes. Letters represent levels of statistical significance between mean AUC values for all constructs (ANOVA, Tukey’s HSD, p<0.05; n=7-8; two independent experiments). **h.** Western blot showing the accumulation of MYC-tagged LNR and single residue mutants in transient expression assays in lettuce cv. Salinas. Untagged LNR is included as a negative control. Detection was performed with α-MYC antibody. Ponceau S staining of blots serves as a loading control. Accumulation was confirmed in an independent experiment.

The predicted structure of the LNR LRR-type ectodomain adopts a solenoid-like shape consisting of 27 LRR repeats, similar to FLS2 and MIK2 (16, 52) (Figure 4d). The interaction between LNR and the nlp24 peptide was mapped to the inner surface of the ectodomain, between LRRs 2 and 19 (Figure 4d, e, Supplementary File 5). Three tested modeling tools (AF3, AF2-Multimer, and Boltz2) predicted that the nlp24 W23 residue inserts into a hydrophobic pocket in the ectodomain surface, interacting with Y522 of LNR via π–π stacking (Figure 4f, Supplementary Figure 14) (53–55). Accordingly, the Y522E substitution in LNR severely disrupted the nlp24-induced ROS burst in transient expression assays (Figure 4g, Supplementary Figure 13). The mutation is predicted to have minimal impact on the LNR protein stability (+0.63 energy units) (56), and its accumulation was confirmed by western blot assays (Figure 4h, Supplementary Figure 13). Overall, the modeling and experimental data suggest that LNR detects the heptapeptide of nlp24 via its LRR-type ectodomain with a crucial role for LNR-Y522 and nlp24-W23 residues.

### LNR belongs to an expanded MIK2 clade that evolved via expansion and contraction in the Asteraceae

Since the Arabidopsis nlp24 receptor, RLP23, does not have homologs in lettuce, we were interested to explore the evolutionary origin of the lettuce LNR receptor. For this, we built a maximum likelihood phylogenetic tree with sequences of LRR-RLK kinase domains from *Arabidopsis* and lettuce. As expected, well-characterized cell surface immune receptors such as *Arabidopsis* FLS2 and EFR fell into the LRR-RLK subfamily XII (11, 12, 17). In contrast, LNR grouped with subfamily XI (Figure 5a, Supplementary Figure 15), also containing *Arabidopsis* CLAVATA1 (CLV1), PEP1 RECEPTOR 1 (PEPR1), and RECEPTOR-LIKE KINASE 7 (RLK7) (57–60). LNR appears to be embedded within a clade of *MIK2-like* genes encoding receptors that recognize microbial and plant SERINE-RICH ENDOGENOUS PEPTIDES (SCOOPs) in *Arabidopsis*, an unknown *Fusarium* elicitor in tomato, and phytocytokines in soybean (Figure 5a, Supplementary Figure 17) (61–65). Members of the MIK2 clade are broadly found in seed plants, and they show strong lineage-specific patterns of duplication and retention (66).

**Figure 5.**
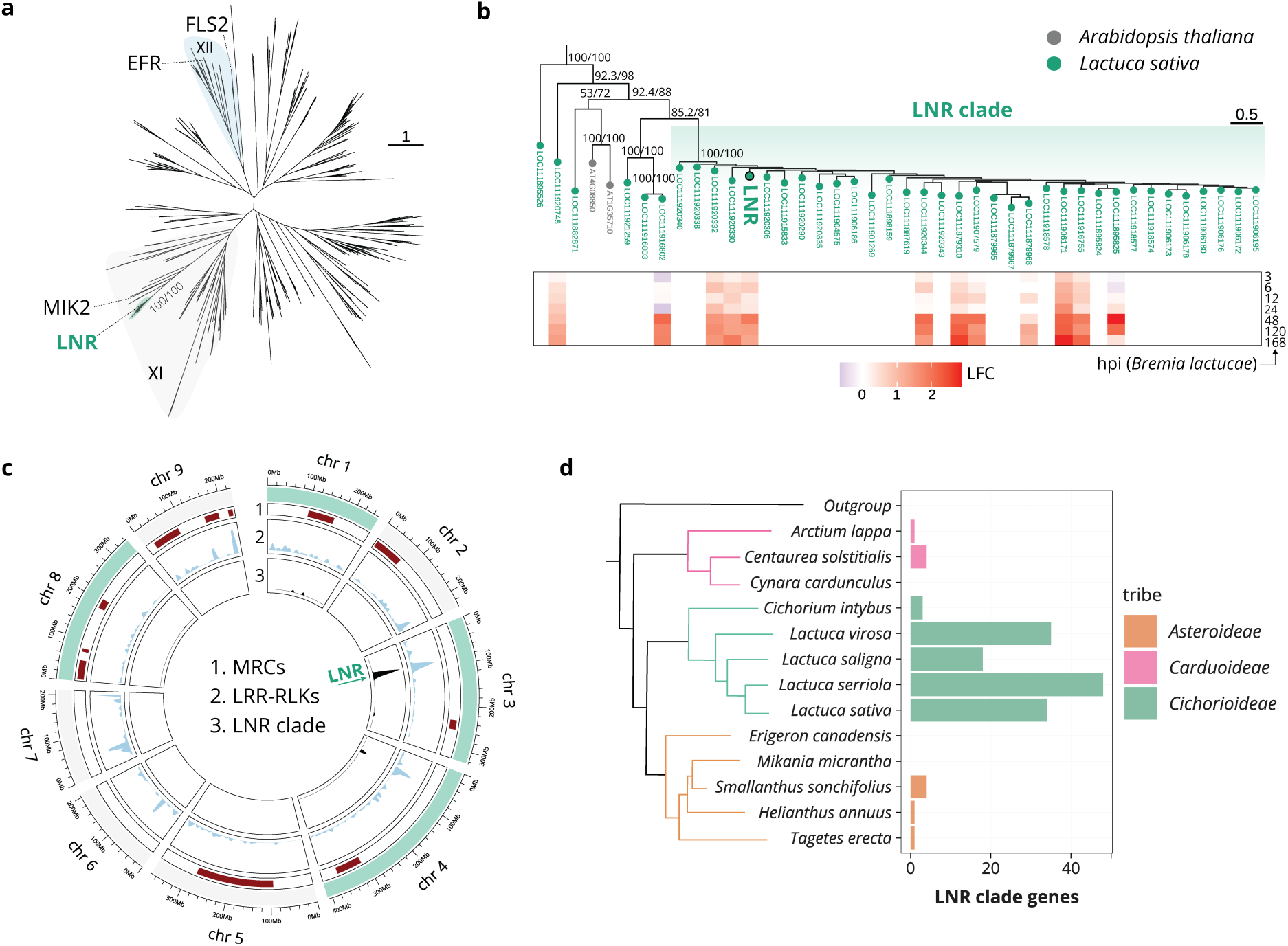
The LNR clade is a close relative of MIK2 and expanded within a locus on chromosome 3 in the lettuce genome. **a.** Unrooted phylogeny of LRR-RLKs in Arabidopsis and *Lactuca sativa*. Subfamilies XI and XII are highlighted (139). Proteins mentioned in the text are labeled, including LNR and its monophyletic group. Ultrafast bootstrap and SH-aLRT support values are included for the LNR clade (1000 replicates). **b.** Zoom-in of the branch leading to the LNR clade. Proteins are colored by species. Ultrafast bootstrap and SH-aLRT support values are indicated for major nodes (1000 replicates). The heatmap shows significant gene expression (log2 fold change) after *Bremia lactucae* infection (3h-7days) (30). **c.** Circos plot of the lettuce reference genome (GCF_002870075.4, v11). Chromosomes with *LNR* clade genes are highlighted in green. Tracks indicate the locations of major resistance clusters (MRCs) (71), *LRR-RLKs*, and genes of the *LNR* clade. **d.** Number of *LNR-like* genes in genomes of the Asteraceae family. Approximate relationships among analyzed species were manually drawn (140). Bar colors indicate the tribe within Asteraceae.

Our synteny analysis identified three genomic regions in the lettuce reference genome with homology to the *Arabidopsis MIK2* gene, consistent with an ancient whole genome triplication in the Asteraceae lineage (67–70), but only one region retained a *MIK2* copy (*LOC111921259*, Figure 5b, Supplementary Figure 16). Although LNR falls within the lettuce MIK2 clade, it belongs to a monophyletic sub-clade with 33 receptor genes in the lettuce reference genome, with no representatives from Arabidopsis or other major angiosperm lineages (Figure 5b, Supplementary Figure 17). *LNR* and *LNR-like* sequences from this sub-clade are spread across chromosomes 1, 3, 4, and 8, located outside both the *MIK2* syntenic regions and the *NLR*-rich major resistance clusters (MRCs) previously described in lettuce (Figure 5c) (71). Notably, most of *LNR-like* genes are clustered near the beginning of chromosome 3 (∼21.4 Mb, 26 of 33 LNR clade sequences, Figure 5c). In the *L. sativa* cv. Ninja genome, but not the Salinas genome, *LNR* is located within this region, ∼0.5 Mb upstream of a local cluster of *LRR-RLK* genes (Figure 3d, Figure 5c). Multiple *LNR-like* genes are upregulated after downy mildew infection by *B. lactucae*, including *LNR* itself (Figure 5b, Supplementary Figure 18) (30). These results show that the locus on chromosome 3 is a hotspot for receptor diversification and evolution in the lettuce genome, likely linked to pathogen detection and plant immunity.

To investigate the origin of the LNR clade, we constructed a maximum likelihood phylogeny with LRR-RLK kinase domains from Asteraceae using representative genomes of major tribes. We also included sequences from the lettuce closest wild relative *L. serriola* and the more distant species *L. saligna* and *L. virosa* (Supplementary Figure 19). The LNR clade formed a well-supported branch (bootstrap value BS=100) with sequences from the *Asteroideae* (e.g., sunflower *Helianthus annuus*), *Carduoidae* (e.g., greater burdock *Arctium lappa*), and *Cichoriodiae* tribes (Supplementary Figure 19). It is striking to see the expansion of *LNR-like* genes in the *Lactuca* genus (on average ∼30 genes per species), while outside the *Lactuca* genus the number of *LNR-like* genes is low (between 0 and 4 genes per species) (Figure 5d). Also, close homologs of the *L. sativa LNR* gene are only found in species of the *Lactuca* genus and not outside of it (Supplementary Figure 19). Taken together, our phylogenetic analysis shows that the LNR-like clade experienced independent losses and expansions after its origin in the common Asteraceae ancestor, resembling the pattern previously described for the broader MIK2 clade across angiosperms (66).

## Discussion

The number of well-characterized cell surface immune receptors from *Arabidopsis* and other species is growing (19–22), however no such receptor has yet been functionally studied in the Asteraceae family, one of the largest plant groups with over 20,000 species and important field and vegetable crops such as sunflower, lettuce, and chicory (72). In this study, we identify LNR as the LRR-RLK mediating the recognition of the nlp24 microbial pattern in cultivated lettuce (*Lactuca sativa*). This was achieved by combining GWAS and presence-absence variation analysis in the diversity panel of 198 lettuce accessions (Figure 3), complementation assays in the non-responsive cv. Salinas and *N. benthamiana* (Figure 4, Supplementary Figure 11, 12), and experimental validation of the structural models for the receptor-peptide interaction (Figure 4, Supplementary Figure 13, 14). LNR belongs to a large *Lactuca*-specific MIK2-like clade of RLKs and not to the RLP23-like protein clade as the nlp24 receptor in *Arabidopsis*, which appears to be Brassica-specific. The different receptors of lettuce and Arabidopsis clearly illustrate how NLP recognition can convergently evolve in each of these species through specific expansion and adaptation of receptor-like sequences.

NLPs show extensive sequence variation, yet a heptapeptide motif (GHRHDWE) is highly conserved across all four NLP classes regardless of their cytotoxicity (26, 36). Our findings demonstrate that this motif constitutes the core epitope of nlp24 recognized by LNR (Figure 1). No receptors were yet found to recognize this NLP motif. In addition to *Arabidopsis* and lettuce, which recognize AIMY and the heptapeptide motifs, respectively, a third NLP recognition specificity has been reported in cucumber. Cucumber responds to the C-terminal 32 residues of *Colletotrichum orbiculare* NLP1 (73), showing that specificities beyond the motifs in nlp24/nlp20 peptides can be observed. Thus, plant species have independently and convergently evolved the ability to recognize different parts of NLPs, which raises a fundamental question about which sequences are likely to be selected as epitopes of immune receptors (74, 75). Gaining knowledge in this direction could facilitate design of functional ligand-receptor pairs.

We found that nlp24 variants lacking the C-terminal heptapeptide (nlp1to11, nlp1to13, nlp1to17, Figure 2d) do not trigger a ROS burst in lettuce, whereas the heptapeptide alone is sufficient for recognition (Figure 2c, d, Supplementary Figure 5). The reduced ROS burst magnitude with the heptapeptide-only fragment, as well as with the variant missing the N-terminal region I (nlp9to24, Figure 2c, d, Supplementary Figure 5), suggests that residues outside the heptapeptide also contribute to full nlp24 recognition, and this complicates linking variation in the heptapeptides of nlp24-like peptides from different microbes to differences in the ROS burst (Figure 2a, b, Supplementary Figure 4). Moreover, our data suggests that residues outside the heptapeptide might not only contribute to the nlp recognition but also take a prominent role in this process when the latter is partially missing. This was observed upon removal of the region I residues with AIMY motif (nlp9to22 and nlp5to22, Figure 2d) in nlp1to22 that abolished the ROS burst (Figure 2c, d, Supplementary Figure 5), suggesting an additive role for the N-terminus in the recognition. However, peptides nlp5to24 and nlp9to24, containing the complete heptapeptide motif, are well recognized. Structural modelling suggests that when the last two residues of the heptapeptide are missing, the phenylalanine in the N-terminal region I of nlp24 takes the space normally occupied by W23 in nlp24 (Supplementary Figure 20). Binding of nlp24 via W23 to LNR is likely more efficient in engaging LNR in defense signaling, since nlp1to22 is a weaker elicitor of immunity than the full-length nlp24 (Figure 2e-g). Thus, LNR is likely a versatile pattern recognition receptor, and this versatility is probably enabled by the flexible π-π stacking between the ligand and the receptor. However, this model of the LNR recognition versatility requires further investigation.

Evolutionarily, LNR belongs to the LRR-RLK subfamily XI (Figure 5a, Supplementary Figure 15), one of the most expanded receptor groups in plants (76). Within subfamily XI, LNR is also part of the MIK2 clade, whose members perceive both endogenous peptides and exogenous immunity triggers (61–66). Within this group, a tomato MIK2-like receptor is involved in the recognition of an unknown *Fusarium oxysporum* elicitor (*Solyc04g074050*, Supplementary Figure 17) (66), while two soybean homologs sense phytocytokines (*Glyma.12G007000* and *Glyma.12G007300*, Supplementary Figure 17) (65). MIK2 (AT4G08850) itself displays dual recognition capacity for SCOOP peptides from *Arabidopsis* and *Fusarium* (Supplementary Figure 17) (61–64). The presence of LNR within this clade emphasizes the versatility of this LRR-RLK group as a scaffold for the recognition of self- and non-self-derived ligands.

Although LNR belongs to the MIK2 clade, *LNR-like* genes form a monophyletic subgroup of 33 receptors in the lettuce reference genome, with no representatives outside of the Asteraceae family (Figure 5b, Supplementary Figure 17). Evolution of the LNR clade in the Asteraceae is shaped by multiple independent losses after its origin in the common ancestor, while the expansion that gave rise to LNR is restricted to *Lactuca* (Figure 5d). This evolutionary profile resembles the evolution of the MIK2 clade across angiosperms, marked by lineage-specific duplications, losses, and retentions (66). Genes of the LNR clade are distributed across four chromosomes, yet only chromosome 3 shows an accumulation of copies organized in local clusters (Figure 3d, Figure 5c). This genomic pattern is consistent with an expansion of this clade via both distal and local tandem duplications, in line with previous observations that RLK- and RLP-encoding genes are often organized in clusters of paralogous genes (76–78). The local accumulation of receptor genes, together with the demonstrated immune function of LNR, suggest that the chromosome 3 locus serves as a hotspot for diversification of the immune receptor repertoire in lettuce.

The evolution of nlp24 perception in the *Lactuca* genus and the mechanisms behind the perception remain to be further explored. Our data suggests that factors beyond *LNR* sequence determine the nlp24 recognition capacity and outputs. On the one hand, nlp24-responsive *L. sativa* accessions display substantial variation in the peptide-induced ROS burst, whereas there is no difference at the *LNR* sequence-level between them (Figure 3a, Supplementary Figure 7d). Furthermore, the wild lettuce relative *L. saligna* does not respond to nlp24 in our ROS assays but encodes a close ortholog of LNR (95.1% protein identity), which is not or very lowly expressed (79). This gene can confer nlp24 responsiveness in transient expression assays in cv. Salinas as determined by the ROS burst assays (Supplementary Figure 11), indicating that the *LNR* presence in the genome alone is not sufficient for the nlp24 recognition. A recent study on legumes showed that the absence of early PTI outputs may not always indicate complete absence of the elicitor recognition and the steady state receptor expression levels could be responsible for this (80). Taken together, these findings indicate that several factors, such as variation in receptor expression, co-receptor(s) availability, or downstream signaling components, could contribute to variation in nlp24 responses in lettuce and remain to be identified.

To conclude, we performed an in-depth characterization of nlp24-triggered immunity in lettuce and identified LNR as the LRR-RLK receptor responsible for its recognition, the first cell surface immune receptor RLK identified and functionally characterized in the Asteraceae family. Its versatile recognition capacity and functionality in Solanaceae provide opportunities for the crop improvement via the receptor design and receptor transfer to other species.

## Materials and Methods

### Plant material and growth conditions

The *L. sativa* diversity panel comprised a total of 198 lines from the Centre for Genetic Resources in the Netherlands (CGN) with available whole-genome resequencing data (46, 47). The complete list of lines used in this study can be found in Supplementary File 6. For experiments, lettuce plants were grown for 14 days on rockwool cubes in a growth chamber at 21 °C under long-day conditions (16h light/8h dark, 100 *μ*mol m^-2^ s^-1^, 70% relative humidity). Hyponex nutrient solution (1x) was provided on the day of sowing and one week after (0.5x). *N. benthamiana* was grown on potting soil for 3 to 4 weeks under the same conditions.

### Peptides

All peptides were synthesized by GenScript. Crude peptides were dissolved in 100% dimethyl sulfoxide (DMSO) to prepare 10 mM stock solutions and diluted in Milli-Q water for their use in experiments. Flg22 100 µM stock solutions (GenScript RP19986) were prepared in Milli-Q water. Alignments of peptide sequences were generated with MAFFT (v7.505, --auto option) (81).

### Identification of nlp24-like peptides from pathogens

NLPs in the *Bremia lactucae* proteome (GCA_004359215.2) were identified using hmmscan against the NPP1 Pfam domain (PF05630) (82). Corresponding nlp24-like sequences were extracted via blastp alignment to the representative nlp24 from *Hyaloperonospora arabidopsidis* NLP3 (28, 83). NLPs in the genome of *Fusarium oxysporum* f. sp. *lactucae* (GCA_045838055.1) were annotated by aligning all proteins containing the NPP1 Pfam domain with miniprot (86), followed by blastp extraction of nlp24-like sequences. Nlp24-like peptides from *Pectobacterium carotovorum, Bacillus subtilis*, and *Phytophthora parasitica* were obtained from previous studies (28, 38).

### ROS measurements

Production of ROS (hydrogen peroxide, H_2_O_2_) was measured in a 96-well plate format with a chemiluminescence-based assay according to (84, 85). The day before the measurements, leaf discs (0 4 mm) were collected from lettuce (first true leaf) or *N. benthamiana* (third and fourth true leaves), washed, and floated overnight in 200 µL Milli-Q water in the dark. To measure the ROS burst, water was replaced with a 250 µL solution containing horseradish peroxidase (HRP, 10 µg/mL, Sigma-Aldrich P8125), L-012 luminol derivative (30 µM, Sigma-Aldrich SML2236), and the corresponding elicitor (2 µM for nlp24 and nlp-like peptides, 0.2 µM for flg22) or a mock (0.02 % DMSO). The luminescence was measured up to 1.5 h on a GloMax96 plate reader (Promega) at 2.5 min intervals. In each experiment, 3-4 leaf discs per treatment were included (technical replicates). Results from independent experiments (biological replicates) were combined for visualization and statistical analysis. Luminescence units over time were log2-transformed and translated into area under the curve (AUC) values with the AUC() function from DescTools R package (86). The normal distribution of residuals and homogeneity of variances were evaluated both visually (Q-Q plots) and statistically with Shapiro-Wilk and Fligner Killeen tests, respectively (α=0.05). Differences among mean values were tested by ANOVA followed by Tukey’s honestly significant difference (HSD) test (α=0.05). Data analysis and visualization was done in R (87). Final figures were edited in Adobe Illustrator.

### ROS measurements in the *L. sativa* diversity panel

Nlp24-triggered ROS production in *L. sativa* was measured in a total of 198 accessions from the diversity panel and four genome reference accessions: cv. Salinas, *L. serriola* CGN25282, *L. saligna* CGN05327, and *L. virosa* CGN04683. ROS production was quantified using the leaf disc-based chemiluminescent assay over 1.5 hours following treatment with 2 µM nlp24 or mock (0.02 % DMSO). Each accession was tested in six independent experiments (biological replicates). For each experiment, leaf discs (n=1 per accession and treatment) were distributed across sixteen 96-well plates, each including positive (cv. Olof) and negative (*L. saligna* CGN05327) controls. Mock- and nlp24-treated discs from each accession were tested on the same plate. Raw luminescence values are available in Supplementary File 4.

Luminescence values per time point were log2-transformed and normalized to the corresponding mock. A total of 622 features were extracted from the ROS burst curves (averaged per treatment) using the Theft R package (88), which provides access to multiple time-series feature sets: Rcatch22 (89, 90), tsfresh (91) and TSFEL (92). AUC values were calculated with AUC() function from DescTools R package (86). Principal component analysis (PCA) on the extracted features was performed with the PCAtools R package (93). The geographical distribution of accessions linked to their nlp24 responsiveness was plotted using the rnaturalearth R package (94), with collection site information obtained from the CGN website.

### MPK phosphorylation assay

Phosphorylation of MPKs was detected via immunoblotting (95). For each sample, nine leaf discs (∅ 4 mm) were collected from lettuce (first leaf) and floated overnight in 2 mL Milli-Q water within a 12-well plate in the dark. The next day, elicitors were directly added to the solution. The final peptide concentration was 2 µM (0.2 µM for flg22) and the mock treatment was 0.02 % DMSO. Upon elicitation, leaf discs were harvested at the indicated time points and frozen in liquid nitrogen. Protein extracts were obtained by grinding the tissue followed by incubation (10 minutes) in extraction buffer (50 mM Tris-HCl pH 7.5, 200 mM NaCl, 1 mM EDTA, 10 % glycerol, 0.1 % Tween20, 1 mM DTT, 1x protease inhibitor (Sigma-Aldrich P9599), 1x phosphatase inhibitor (Sigma-Aldrich P2850)). The supernatants were collected by centrifugation at 18,000g for 30 minutes and boiled in 1x Laemmli buffer at 95 °C for 10 minutes. Denatured protein extracts were resolved in 4-12% SDS-PAGE (ThermoFisher NW04120BOX) and transferred (semi-dry, Bio-Rad 1704150) to nitrocellulose membranes (Bio-Rad 1704158). Ponceau S staining of the membranes served as a loading control (Sigma-Aldrich 09276). MPK3 and MPK6 protein levels were detected with α-MPK3/6 antibodies (Sigma-Aldrich M8318 and A7104, rabbit, 1:2500 dilution), and phosphorylated MPKs with α-p44/42-ERK antibody (Cell-Signaling Technology 9101, rabbit, 1:2500 dilution). An HRP-linked secondary antibody was used (Cell-Signaling Technology 7074, 1:2500 dilution). For signal detection, the chemiluminescent assays Clarity and Clarity max (Bio-Rad 1705061 and 1705062) and the Azure 280 imaging system (Azure biosystems) were employed. Lettuce MPK3 and MPK6 and their molecular weights were described in a previous study (96). Final figures were prepared in Adobe Illustrator.

### Molecular cloning

Expression constructs for membrane receptors were generated via Golden Gate cloning and modules available in the MoClo Plant Parts kits I and II (97, 98). To create level 0 vectors with our genes of interest, sequences without the stop codon were first amplified from genomic DNA of cv. Ninja (*LNR, LSAT002_V1C300114880*), cv. Salinas (*LNRparalog*, *LOC111920335*), *L. saligna* CGN05327 (*LNRsaligna*, *LSALG_LOCUS16100*), or *Arabidopsis* Col-0 (*RLP23*, *AT2G32680*). DNA extractions were performed with the MagMax^TM^ Plant DNA isolation kit (ThermoFisher A47156). To enable Golden Gate cloning, gene sequences were domesticated by removing internal Type IIs restriction sites. This was achieved by introducing synonymous mutations at wobble bases of the affected codons during PCR amplification. The required single nucleotide changes were added to the 4-bp primer overhangs between consecutive fragments, ensuring their introduction into the sequence during PCR. All PCRs were done with Phusion (Thermo Fisher F530) or PrimeSTAR GXL DNA polymerase (Takara Bio R050A). Gene fragments that could not be amplified by PCR were synthesized as gBlocks (Integrated DNA Technologies). Genes were assembled and cloned into the level 0 vector pAGM1287 in a single Golden Gate reaction. Resulting level 0 vectors with the receptor-coding sequences were combined with level 0 vectors containing the lettuce polyubiquitin promoter and 5’ UTR (pICH41295-LsUBIpro), a stop codon (pAGM1301-stop), and the nos terminator with 3’ UTR (pICH41421) into the level 1 backbone pICH47732. The module with a stop codon was replaced by level 0 vectors for mEGFP (pJOG176) or 4x-MYC (pICSL50010) to create C-terminally tagged units. Level 2 assemblies in pAGM4673 backbone included the reconstructed transcriptional unit at position 1 and the coding sequence for *NPTII* kanamycin resistance cassette (pICH47742-NOSpro-NPTII-OCSterm) at position 2. Empty positions at level 2 were filled in with an end-linker (pICH41744). Modules with the parsley polyubiquitin promoter (pICH41233-pcUBIpro), the tobacco mosaic virus 5’ UTR (pICH41402), and GFP (pICSL50008) replacing their counterparts were generated for *LNR* and employed for ectopic transient expression in *N. benthamiana* during *P. capsici* infection assays. All vectors were verified by Sanger sequencing. The complete list of primers used for cloning is provided in Supplementary File 7.

### Site-directed mutagenesis

Substitutions in the *LNR* sequence (Y522E, D958N) were introduced using the QuikChange Site-Directed Mutagenesis protocol (Agilent 200518) and the PrimeSTAR GXL DNA polymerase (Takara Bio R050A). Overlapping primers with the desired single nucleotide changes were designed according to the protocol recommendations (Supplementary File 7). Level 0 constructs encoding Y522E and D958N variants of LNR were first cloned into level 1 vector pICH47732 to create untagged and C-terminally tagged modules (mEGFP, 4x-MYC), and then into the level2 pAGM4673 backbone together with the *NPTII* kanamycin resistance cassette via Golden Gate reactions. The predicted effect of Y522E mutation in the AlphaFold3-modeled LNR ectodomain structure (measured as a change in energy units) was determined using the MutationExplorer server (99).

### RNA extraction, library preparation, and sequencing

The first true leaves of lettuce cv. Olof plants were infiltrated with 0.5 µM nlp24 or mock (Milli-Q water) using a needleless syringe. Per sample, three leaf discs (0 10 mm) from three separate plants were pooled together and frozen in liquid nitrogen 1h post-infiltration. Total RNA was isolated with the MagMAX^TM^ plant RNA isolation kit (Thermo-Fisher A33899), and mRNA libraries were prepared via poly(A) enrichment (stranded, illumina TruSeq kit). Sequencing was performed on a NovaSeq 500 platform (2×150 bp reads, ∼400 million clusters), yielding an average of 55.2 million reads per sample. Library preparation and sequencing were conducted at the Utrecht Sequencing Facility (USEQ).

### RNA-seq data analysis

Gene counts were obtained with the RNASeq-NF pipeline (https://github.com/UMCUGenetics/RNASeq-NF/). Briefly, adapter sequences and low-quality ends of the reads were filtered out with TrimGalore (v0.6.5) (100), and rRNA-derived reads were removed with SortMeRNA (v4.3.3) (101). Filtered reads were aligned to the lettuce cv. Salinas reference genome (GCF_002870075.4, v11) with STAR (v2.7.3a) (102), and gene counts were obtained with featureCounts (subread R package v2.0.0) (103, 104), based on the published gene annotation combining RefSeq and non-overlapping genes from the original NCBI submission (105). Processing of raw reads into gene count tables was conducted at USEQ.

Downstream analysis of gene counts was performed with the DESeq2 R package (106). Lowly expressed genes were first discarded; only those with more than 10 counts in at least 2 samples were retained. Differential gene expression analysis between nlp24- and mock-treated samples was done with the Wald test implemented in DESeq2. Genes with |LFC|>=1 and adjusted p-value<=0.05 (Benjamini-Hochber) were considered as differentially expressed (Supplementary File 1).

To test whether nlp24-upregulated genes in lettuce significantly overlap with the set of commonly upregulated genes during PTI in *Arabidopsis* (43), a one-sided Fisher’s exact test was employed (fisher.test() R function, alternative=’greater’) (87). One-to-one homology relationships between genes in these species were determined with MMseqs2 (105, 107), and all expressed lettuce genes were used as the background. On the volcano plot, differentially expressed lettuce genes were labeled as ‘immunity-related’ when the *Arabidopsis* homolog was linked to a defense-related Gene Ontology (GO) term (descendants of GO:0002376 -immune system process-, GO:0006952 -defense response-, and GO: 0009607 -response to biotic stimulus-).

GO enrichment on differentially expressed genes was performed with the enrichGO function (with parameters pAdjustMethod = "BH", qvalueCutoff = 0.05) implemented in the clusterProfiler R package (108), with all expressed lettuce genes as the background. The homologs in *Arabidopsis* for the lettuce genes as determined by MMseqs2 were used for this analysis (105, 107). GO terms with an adjusted p-value <= 0.05 (Benjamini-Hochberg) were considered significant (Supplementary File 2). GO enrichment was conducted separately for upregulated and downregulated genes, with no enriched terms in the latter.

To analyze the enrichment of TF binding motifs in the promoters of lettuce genes differentially expressed by nlp24, 1500 bp sequences upstream of the coding sequence were extracted from the lettuce reference genome (GCF_002870075.4, v11). A simple enrichment analysis (SEA) was performed to test for enrichment of *Arabidopsis* TF binding motifs with the promoters of all expressed genes as control (109). Motifs from the DAP-seq and protein-binding microarray (PBM) Arabidopsis databases were used for the search (110, 111). SEA is part of the MEME (v5.5.8) suite of tools (109).

### Quantitative PCR

Defense-related marker gene expression analysis upon peptide elicitation was performed via quantitative PCR (qPCR). For each sample, nine leaf discs (∅ 4 mm) were collected from lettuce cv. Olof (first leaf) and floated overnight in 2 mL Milli-Q water within a 12-well plate. The next day, elicitors were gently added to the solution up to the desired concentrations: 2 µM nlp24, 2 µM nlp1to22, 0.2 µM flg22, or mock (0.02 % DMSO). Upon elicitation, leaf discs were harvested at the indicated time points and frozen in liquid nitrogen. Total RNA was isolated with the MagMAX^TM^ plant RNA isolation kit (Thermo-Fisher A33899), which includes a DNaseI treatment for genomic DNA removal. Total RNA (1 µg) was used for cDNA synthesis with RevertAid H Minus Reverse Transcriptase (Thermo-Fisher EP0452) and odT18VN primer (Integrated DNA Technologies). The newly synthesized cDNA was diluted and used as input for qPCR with the iTaq Universal SYBR Green mix (Bio-Rad 172-5125) and gene-specific primers (0.5 µM each). The CFX Opus 384 real-time system (Bio-Rad 12011452) was employed with the standard qPCR protocol. Primers for *LOC111886289* (*EIX2*), *LOC111876388* (*WRKY30*), and *LOC111884022* (*WRKY55*) were designed with the NCBI primer-blast tool (112) and validated by performing a standard curve using a serial dilution of cDNA template. The expression of each gene was normalized to *LOC111882438* (*ACTIN7*). In the statistical analysis, results from four separate experiments (biological replicates) were combined. Each sample was tested three times (technical replicates). The normal distribution of residuals and homogeneity of variances were evaluated both visually (Q-Q plots) and statistically with Shapiro-Wilk and Fligner Killeen tests, respectively (α=0.05). Differences among mean values were tested by ANOVA followed by Tukey’s honestly significant difference (HSD) test (α=0.05).

### Infection assays with *Bremia lactucae* in lettuce after peptide treatment

The first true leaves of lettuce cv. Olof plants were fully infiltrated with 2 µM nlp24, 2 µM nlp1to22, or mock (0.02% DMSO) using a needleless syringe. After 24 hours, plants were spray-inoculated with a spore suspension of *Bremia lactucae* isolate Bl:21 (100 conidiospores/µL). Inoculated plants were kept inside covered trays and placed in a growth chamber at 16 °C under short-day conditions (9h light/16h dark, 100 *μ*mol/m^2^/s). The first and second true leaves were collected separately at 13 dpi to quantify local and systemic effects of peptide pre-infiltration on *B. lactucae* sporulation. In the statistical analysis, results from three independent experiments were combined. The normal distribution of residuals and homogeneity of variances were evaluated both visually (Q-Q plots) and statistically with Shapiro-Wilk and Fligner Killeen tests, respectively (α=0.05). Differences among mean values were tested by ANOVA followed by Tukey’s honestly significant difference (HSD) test (α=0.05).

### Segregation analysis

An F_2_ population was generated from the cross between cv. Crna (CGN16263, responsive to nlp24) and cv. Salinas (CGN25281, non-responsive to nlp24). A total of 194 F_2_ individuals were phenotyped for responsiveness to nlp24 at the ROS production level. Segregation data from the F_2_ individuals were tested against a 3:1 Mendelian ratio with a chi-square (χ^2^) test (chisq.test() R function) (87).

### Genome-wide association studies (GWAS)

GWAS for nlp24-triggered ROS burst in the *L. sativa* diversity panel of 198 accessions was performed using the area under the ROS curve as the phenotypic value. This feature was among the top contributors to PC1 in the previously described PCA (Supplementary Figure 7). The SNP matrix for this population was obtained from a previous study (48), in which variants were called by aligning publicly available resequencing data to the cv. Salinas reference genome (GCF_002870075.3, v8). A total of 2,491,009 variants were annotated. GWAS was conducted with the lme4QTL R package (49) using scripts from an earlier publication (50). The SNP covariance matrix was used to account for population structure (48). Associations exceeding the Bonferroni-adjusted significance threshold (–log_10_ (p-value) > 7.69) were considered significant. All significant 5,031 SNPs are listed in Supplementary File 8, including their estimated positions in the latest cv. Salinas reference genome assembly (GCF_002870075.4, v11) based on gene proximity.

### Visualization of genomic loci

The locus on chromosome 3 significantly associated with nlp24 perception (77,980,663-88,200,496) was extracted from the latest cv. Salinas reference genome assembly (GCF_002870075.4, v11) with SeqKit (113). The homologous region in cv. Ninja chromosome 3 (77,128,571-87,011,805) was identified by MUMmer-based alignment (114) (‘nucmer –t 24 -l 500 -g 1000’). Gene annotations for cv. Salinas were extracted from the published annotation (105), and for cv. Ninja from the MAKER annotation submitted to NCBI (*assembly to be released in GeneBank*). Genes were classified as RLK- or RLP-encoding with custom scripts (*see ‘RLK and RLP annotation and classification’*). Local sequence homologies (>= 20 Kbp) were identified via Minimap2 (command ‘minimap2 -X -N 50 -p 0.1 -c’) (115). Transposable elements (TEs) were annotated *de novo* by first generating consensus sequences per TE family with RepeatModeler (116), followed by masking with RepeatMasker (117). Genomic loci and their annotations were visualized with the gggenomes R package (118), and the final figure was edited in Adobe Illustrator.

### RLK and RLP annotation and classification

All genomes analyzed in this study (Supplementary File 9) were searched for RLK- and RLP-encoding genes following the same procedure. TMbed (v1.0.0) (119) was employed to detect signal peptides and transmembrane domains in all proteins per genome (single isoform per gene). Sequences with at least a transmembrane region were scanned against the Pfam-A protein family database with the pfam_scan.pl script (options -e_seq 0.1 -e_dom 0.1) (120). Proteins containing a PRR-related extracellular domain, a transmembrane segment, and an intracellular kinase domain were classified as RLKs. Proteins meeting these criteria but missing the kinase domain were annotated as RLPs. Classification of RLKs and RLPs into different classes was done according to the ectodomain type. The domains used for RLK and RLP annotation and classification are listed in Supplementary File 10. The number of annotated LRR-RLKs was verified by comparison with annotations from other studies (76, 121) (Supplementary File 11).

### LNR coverage from whole-genome resequencing data

Whole-genome resequencing data for the 198 accessions in the *L. sativa* diversity panel were downloaded from the CNGB Nucleotide Sequence Archive (CNSA, accession number CNP0000335) and the European Nucleotide Archive (ENA, accession number PRJEB63589). Adapter sequences and low-quality ends of the reads were trimmed with TrimGalore (v0.6.5) (100), and reads were then aligned to the cv. Ninja genome with bwa-mem2 (122, 123). Read group information was preserved in the alignment. *LNR* coverage (defined as the proportion of its length covered by at least one read) was calculated with the ‘coverage’ function in bedtools (v2.31.1) (124). The presence of distinct *LNR* haplotypes across the *L. sativa* accessions was tested by variant calling. Coordinate-sorted, duplicate-marked alignments were processed with FreeBayes (v1.3.9, option ‘min-mapping-quality 20’) (125), but no variants were detected.

### Transient expression assays

The same procedure was followed for transient expression assays in lettuce cv. Salinas and *N. benthamiana*. Expression constructs for *LNR*, single residue variants of *LNR* (Y522E and D958N), *RLP23*, *LOC111920335* (*LNR* paralog in cv. Salinas, negative control), and *Lsal_1_v1_gn_3_00002013* (*LNR* ortholog in *L. saligna* CGN05327), with or without a C-terminal tag (mEGFP or 4xMyc), were electroporated into *Agrobacterium tumefaciens* AGL1 supplemented with a constitutive *virG* mutant gene for enhanced T-DNA transfer efficiency (126–128). For transient expression, *A. tumefaciens* with the indicated plasmids were grown on LB agar plates with antibiotics for 2-3 days, followed by overnight growth on fresh plates. Each strain was then mixed with *A. tumefaciens* C58C1 expressing *p19* to a final OD_600_ of 0.2. Co-cultures were prepared in induction buffer (10 mM MgCl_2_, 10 mM MES pH 5.6, 150 µM acetosyringone) and incubated for 2-3 hours in the dark at room temperature. The first true leaves of lettuce cv. Salinas and the third to fourth true leaves of *N. benthamiana* were infiltrated with a needleless syringe. Leaf discs for ROS detection and immunoblot analysis of expressed receptor constructs were collected 2 days post-*Agrobacterium* infiltration.

### Immunoblot analysis of membrane receptors

To assess the accumulation of transiently expressed receptor constructs in lettuce, two 10-mm leaf discs were harvested at 2 days post-*Agrobacterium* infiltration and frozen in liquid nitrogen. Protein extracts were prepared by grinding the tissue and incubating it for 2 h in extraction buffer (25 mM Tris-HCl pH 8, 150 mM NaCl, 1% IGEPAL (Sigma-Aldrich I8896), 0.5% SDS, 1x protease inhibitor (Sigma-Aldrich P9599), 1x phosphatase inhibitor (Sigma-Aldrich P2850)). The supernatants were collected by centrifugation at 18,000g for 30 minutes and boiled in 1x Laemmli buffer at 95 °C for 10 minutes. Denatured protein extracts were resolved in 4-12% SDS-PAGE (ThermoFisher NW04120BOX) and transferred (semi-dry, Bio-Rad 1704150) to nitrocellulose membranes (Bio-Rad 1704158). Ponceau S staining of the membranes served as a loading control (Sigma-Aldrich 09276). Membranes were incubated with primary antibodies α-GFP (Cell-Signaling Technology 2956, rabbit, 1:2500 dilution) or α-MYC (Cell-Signaling Technology 2278, rabbit, 1:2500 dilution), and an HRP-linked secondary antibody was used (Cell-Signaling Technology 7074, 1:2500 dilution). For signal detection, the chemiluminescent assays Clarity and Clarity Max (Bio-Rad 1705061 and 1705062) and the Azure 280 imaging system (Azure biosystems) were employed. Final figures were prepared in Adobe Illustrator.

### *Phytophthora capsici* infection assays in *N. benthamiana*

*Phytophthora capsici* isolate LT3112 was cultured on V8 agar plates (20% V8 juice, 0.02% CaCO_3_, 1.5 % agar) at 21°C in a growth chamber under long-day conditions (16h light/8h dark). Infection assays were performed on *N. benthamiana* detached leaves transiently expressing an empty vector (pAGM4673) or constructs with *LNR*, or *LNR-GFP*. Three days after *Agrobacterium* infiltration, the third and fourth true leaves were detached and placed in square Petri dishes on three water-saturated filter papers (80 mm). Leaves were then inoculated with *P. capsici* mycelium plugs from 5-day-old cultures, and sealed plates were maintained in a growth cabinet (21 °C, 16h light/8h dark). Disease symptoms were evaluated two days after infection by measuring lesion size from RGB images using Fiji (129). Per experiment, 14-16 lesions were analyzed (technical replicates), except for the second experiment (8 lesions). In the statistical analysis of lesion size measurements, results from four independent experiments were combined. The normal distribution of residuals and homogeneity of variances were evaluated both visually (Q-Q plots) and statistically with Shapiro-Wilk and Fligner Killeen tests, respectively (α=0.05). Differences among mean values were tested by ANOVA followed by Tukey’s honestly significant difference (HSD) test (α=0.05).

### Structural modeling of the receptor-peptide interaction

The interaction between the LNR ectodomain (residues 34-767) and nlp24 was modeled using three different algorithms: AlphaFold2 (55), AlphaFold3 (53), and Boltz2 (54). AlphaFold2 (v2.3.2-1) and Boltz2 (v2.2.1) were run on a local HPC equipped with CUDA-enabled GPUs. AlphaFold3 predictions were obtained in the online server (https://alphafoldserver.com/). The top scored prediction per algorithm was used for visualizations. Atomic-level interactions in the AlphaFold3-modeled receptor-peptide complex were profiled with the PLIP server (https://plip-tool.biotec.tu-dresden.de) (130). Contacts are listed in Supplementary File 5. The interaction between the LNR ectodomain and nlp1to22 was modeled in the AlphaFold3 server. Visualizations were done in ChimeraX (v1.6.1).

### Phylogenetic analysis and visualization

The proteomes of all species analyzed in this study were screened for RLK and RLP sequences using the domain-based annotation described above (Supplementary File 11). For reconstruction of LRR-RLK phylogenies, intracellular kinase domains were extracted with SeqKit (113) and subsequently aligned with MAFFT (v7.505, l-ins-i or fft-ns-2 algorithm) (81). Alignments were trimmed with ClipKit (v2.4.1, -m kpic-smart-gap) (131) and used as input for maximum-likelihood phylogeny inference with IQ-TREE (v3.0.1, -m MFP -B 1000 -bnni -alrt 1000 -T auto) (132). The best-fit substitution model for the data was determined by ModelFinder (133). LNR was added into cv. Salinas protein sequences during tree reconstruction. Resulting trees were visualized with the ggtree R package (134).

The phylogeny of LRR-RLKs in *Arabidopsis* and lettuce was annotated with RNA-seq expression data from lettuce infected with *Bremia lactucae* SF5 isolate (30). Sequencing reads were downloaded from ENA (accession PRJNA523226) and processed on a local HPC. Adapters and low-quality ends of the reads were filtered out with TrimGalore (v0.6.5) (100), and rRNA-derived reads were removed with SortMeRNA (v4.3.3) (101). Expression levels were quantified against the cv. Salinas transcriptome (complemented with the *LNR* transcript) using Salmon (v1.9.0, salmon quant, -l SR --seqBias -- discardOrphansQuasi) (135). Transcript-to-gene level translation of expression values was performed with the tximport R package (136), and differential expression was assessed with DESeq2 (106) by comparing each infection time point against 0 hpi.

### Visualization of genomic features in the lettuce reference genome

Locations in the cv. Salinas lettuce reference genome of the major resistance clusters (MRCs) (71), LRR-RLKs, and LRR-RLKs belonging to the LNR clade were visualized with the circlize R package (137).

### Synteny analysis

Synteny between the *Arabidopsis* locus on chromosome 4 containing *MIK2* (chr4: 5,136,479-6,140,952; gene location extended by 500 Kbp) and the cv. Salinas genome (GCF_002870075.4, v11) was assessed with MCScanX (138). Proteins in the *AtMIK2* locus were aligned against single-isoform lettuce proteins using blastp (v2.15.0+, -evalue 1e-10 -max_hsps 1 -max_target_seqs 5) (83). Results were provided as input to MCScanX with default parameters. Synteny blocks were visualized with circlize R package (137).

### Data availability

The raw RNA sequencing data were uploaded to Short Nucleotide Archive at NCBI (PRJNA1354397; https://dataview.ncbi.nlm.nih.gov/object/PRJNA1354397?reviewer=hue301ceenm70ipf2ksbph0gq4). The original code used in this study will be available under https://github.com/inibb/nlp24_receptor_lettuce. Other data is available upon request to the corresponding authors.

## Acknowledgements

We thank Michael Seidl for helpful discussions during the development of this project. We are also grateful to Alexander Kozik for providing sequences from the *cv.* Ninja genome assembly, Dirk-Jan van Workum for contributions to the discussions on the RLK/RLP classification code, and Flip Mulder for processing the raw RNA-seq data into gene count tables.

This publication is part of the LettuceKnow project (with project number 2.1B of the research Perspective Program P19-17 which is (partly) financed by the Dutch Research Council (NWO) and the breeding companies BASF, Bejo Zaden B.V., Limagrain, Enza Zaden Research & Development B.V., Rijk Zwaan Breeding B.V., Syngenta Seeds B.V., and Takii and Company Ltd. Research reported in this publication was supported in part by award # 0000000001 from the Foundation for Food and Agriculture Research to RWM. The content of this publication is solely the responsibility of the authors and does not necessarily represent the official views of the Foundation for Food and Agriculture Research. IB, GVDA, and DL are inventors on a patent application describing the utilization of the LETTUCE nlp24 RECEPTOR in the disease resistance improvement. These and other authors declare no other competing interests.

## Author contributions

Conceptualization – IB, DL, GVDA; Methodology – IB, DL, GVDA, BS; Software – IB, SS, SM, BS; Validation – DL, GVDA; Formal analysis – IB, SS, SM; Investigation – IB, SM, SAL, MA, SS, MP, SB, AVB, PS, SP, GVDA, BS; Resources – MA, RM; Data curation – IB, MA, SM, DL; Writing, original draft and preparation – IB, DL, GVDA; Writing, review and editing – IB, BS, RM, DL, GVDA; Visualization – IB; Supervision – DL, GVDA, BS; Project administration – RM, DL, GVDA; Funding acquisition – RM, DL, GVDA.

