## Supplementary Information for "A lettuce receptor-like kinase recognizes the highly conserved heptapeptide motif within microbial NEP1-like proteins"

**This PDF contains:**

Figures S1 to S20

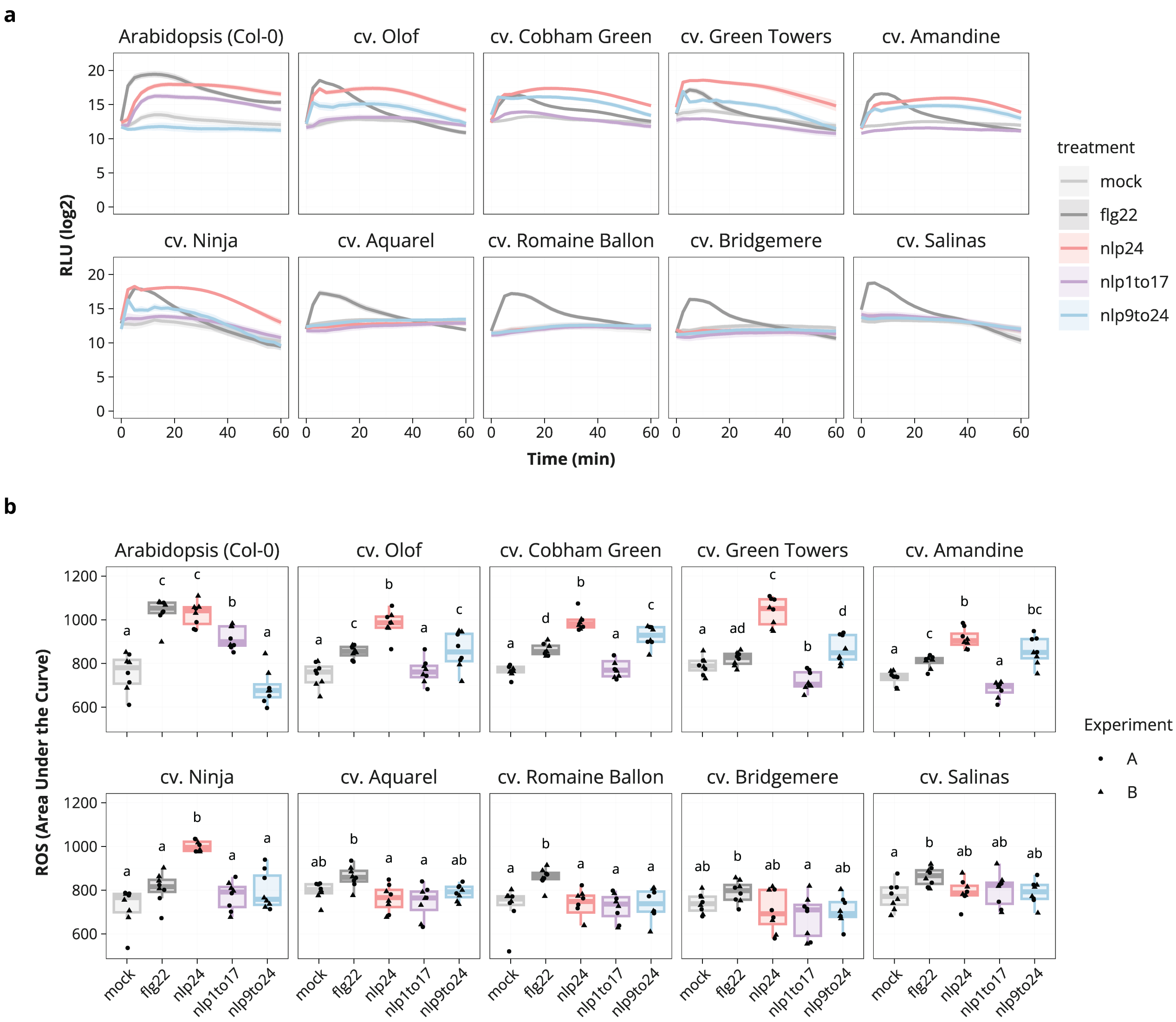


**Supplementary Figure 1.** Reactive oxygen species (ROS) production in *Arabidopsis* Col-0 and nine lettuce cultivars after treatment with flg22 (0.2 μM), nlp24 (2 μM), nlp1to17 (2 μM), nlp9to24 (2 μM), or mock (0.02 % DMSO). **a.** ROS burst over 1 hour. Mean and standard error per time point is shown (n=8, two independent experiments). **b.** ROS burst translated to area under the curve (AUC) values. Letters represent levels of statistical significance between mean AUC values (ANOVA, Tukey's HSD, $p<0.05$).

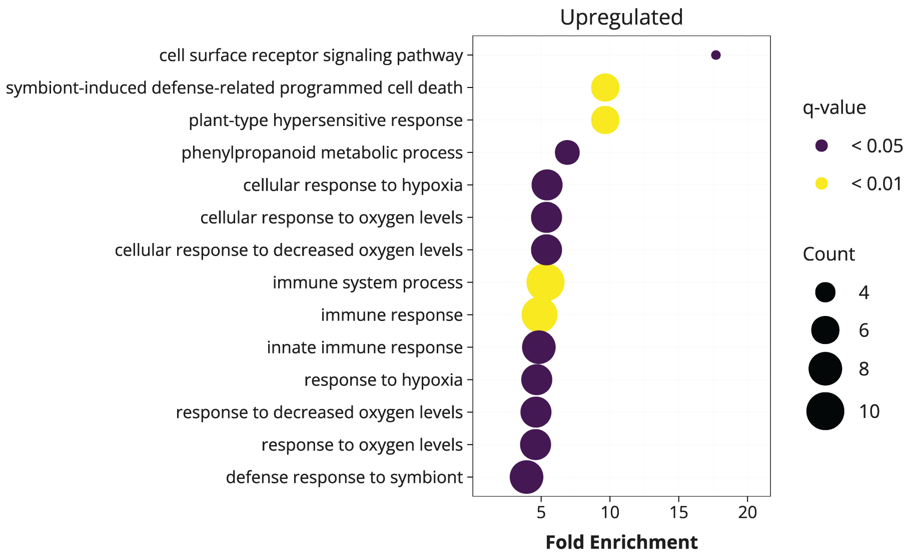


**Supplementary Figure 2**. Gene ontology (GO) term enrichment analysis of nlp24-upregulated genes in lettuce cv. Olof (1 h after infiltration) based on closest *Arabidopsis* homologs. The analysis was done with enrichGO function from clusterProfiler R package. P-values were adjusted using the Benjamini–Hochberg method (q-value cutoff 0.05). Term size represents the number of genes in the GO term, and color corresponds to the adjusted p-value.

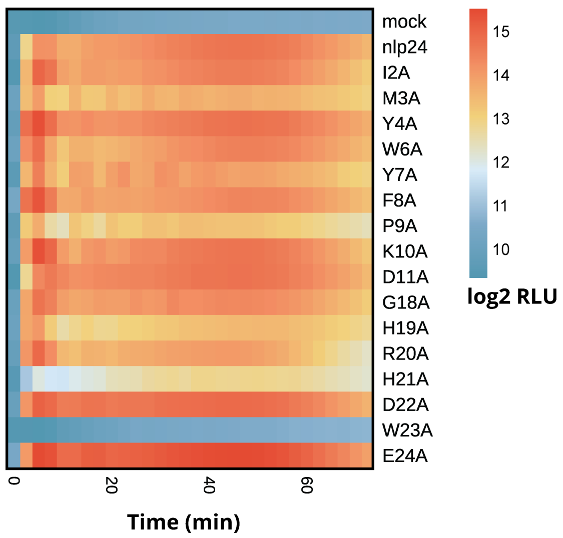


**Supplementary Figure 3.** ROS burst in lettuce cv. Olof treated with nlp24 (2 µM), alanine substitution variants (2 µM), or mock (0.02 % DMSO). Mean log2 RLU values are shown on the heatmap; the experiment was repeated three times.

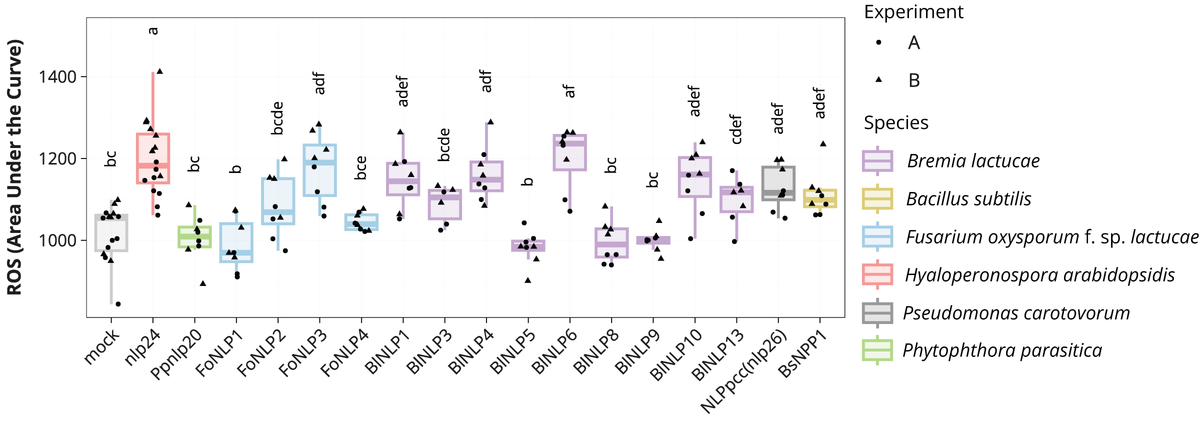


**Supplementary Figure 4**. ROS burst in lettuce cv. Olof after treatment with nlp24-like peptides from *Fusarium oxysporum* f. sp. *lactucae* (Fo), *Bremia lactucae* (Bl), *Pectobacterium carotovorum* (Pcc), and *Bacillus subtilis* (Bs) (2 µM), nlp20 from *Phytophthora parasitica* (Pp) (2 µM), or with mock (0.02 % DMSO). The ROS burst was measured over 82.5 minutes and translated to AUC values. Letters represent statistically significant differences (ANOVA, Tukey's HSD, $p<0.05$; n=15 for mock, n=16 for nlp24, n=6-8 for other peptides; two independent experiments).

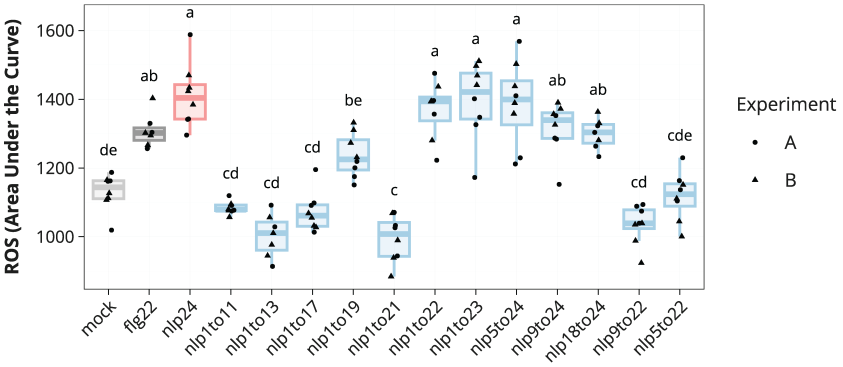


**Supplementary Figure 5**. ROS burst in lettuce cv. Olof after treatment with nlp24 and truncated variants (2 µM). The ROS burst was measured over 87.5 minutes and translated to AUC values. Letters represent statistically significant differences (ANOVA, Tukey's HSD, $p<0.05$; n=8 for mock; two independent experiments).

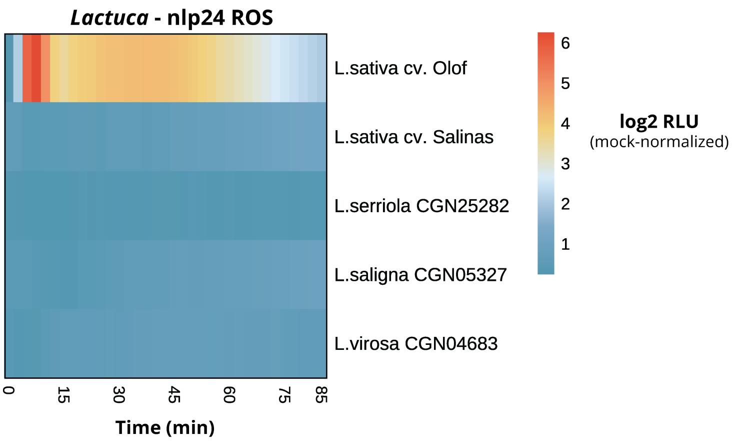


**Supplementary Figure 6**. ROS production in *L. sativa* cv. Olof, cv. Salinas, *L. serriola* CGN25282, *L. saligna* CGN05327, and *L.virosa* CGN04683, after nlp24 treatment (2 µM, normalized to mock). Mean values per time point are shown (n=87 for *L. saligna* CGN05327, n=21 for *L. virosa* CGN04683, and n=22 for the rest). Combined results from six experiments are shown.

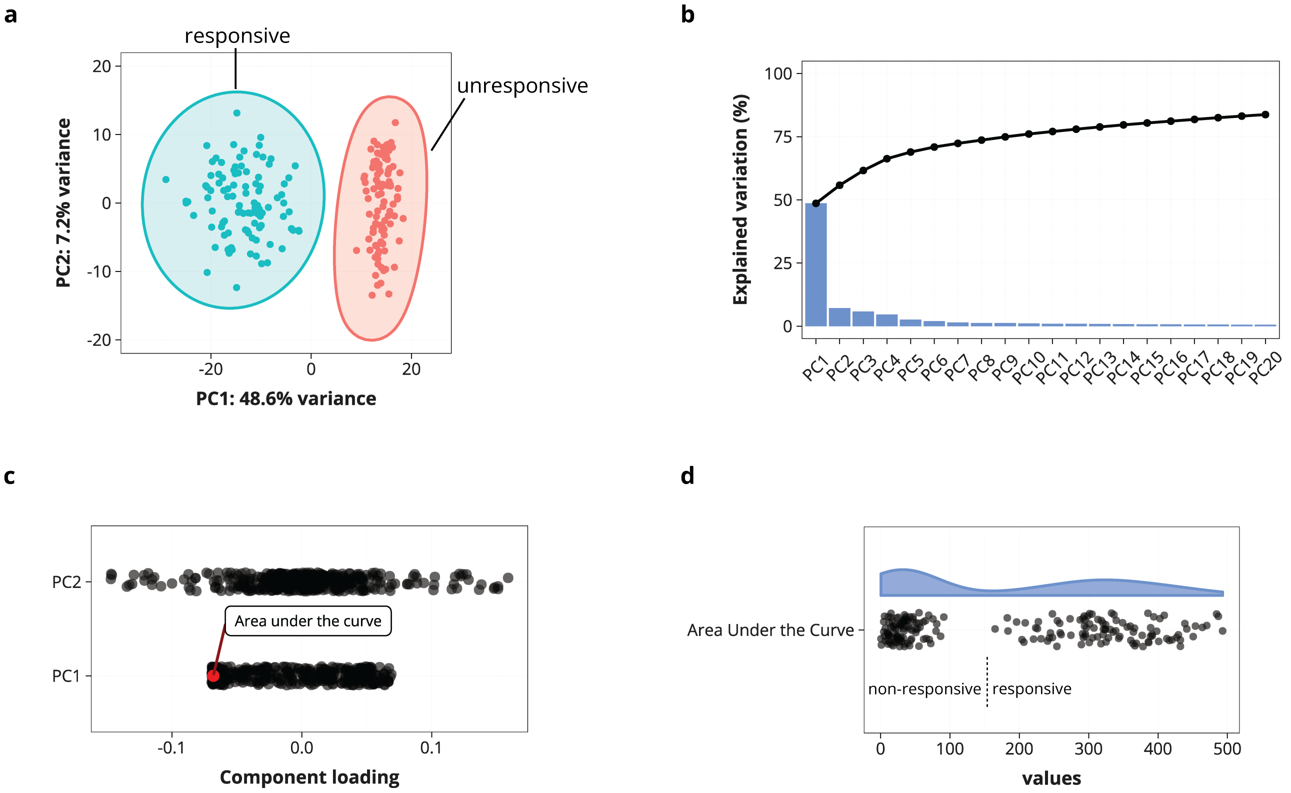


**Supplementary Figure 7.** PCA analysis of features extracted from nlp24-triggered ROS curves in 198 *L. sativa* accessions. ROS production values were normalized to mock prior to feature extraction. **a.** Biplot of 474 features extracted from ROS production curves over time. X- and Y-axis contain the first two principal components (PC1, 48.6 % variance; PC2, 7.2 % variance). Circles surrounding points are colored based on nlp24 responsiveness. **b.** Barplot showing the proportion of variance explained by each PC. **c.** Component loadings for PC1 and PC2. Each dot represents the contribution of a feature to the corresponding PC. **d.** AUC values for the 198 *L. sativa* accessions after nlp24 exposure.

a

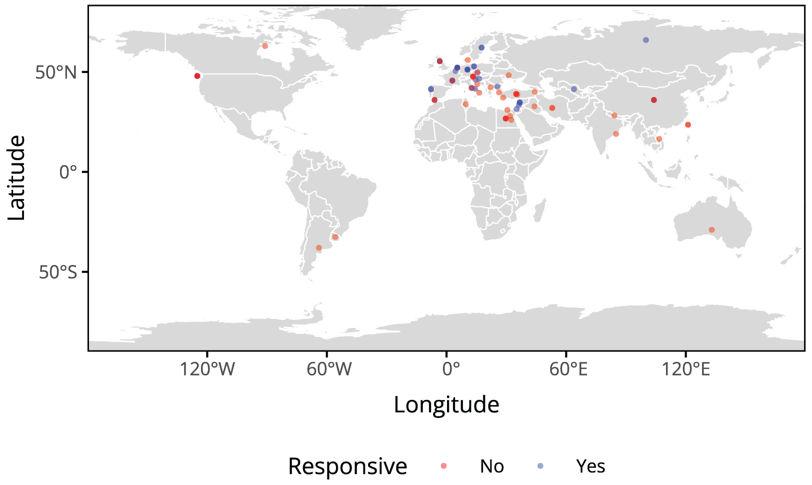


b

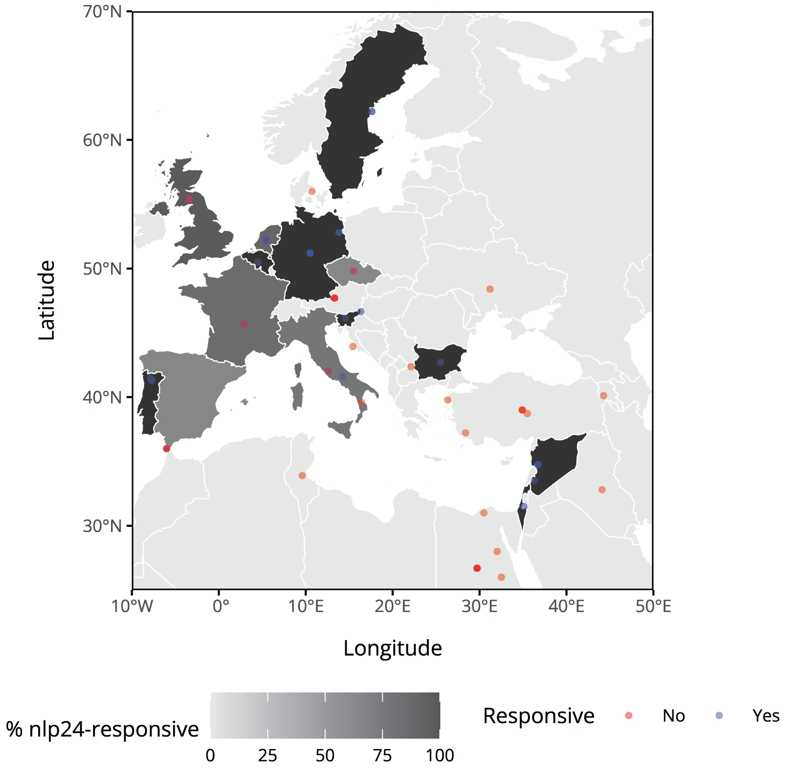


**Supplementary Figure 8.** Map of geographic distribution of nlp24 responsiveness across *L. sativa* accessions. **a.** Distribution across all countries. **b.** Zoom into the region where most accessions originate from.

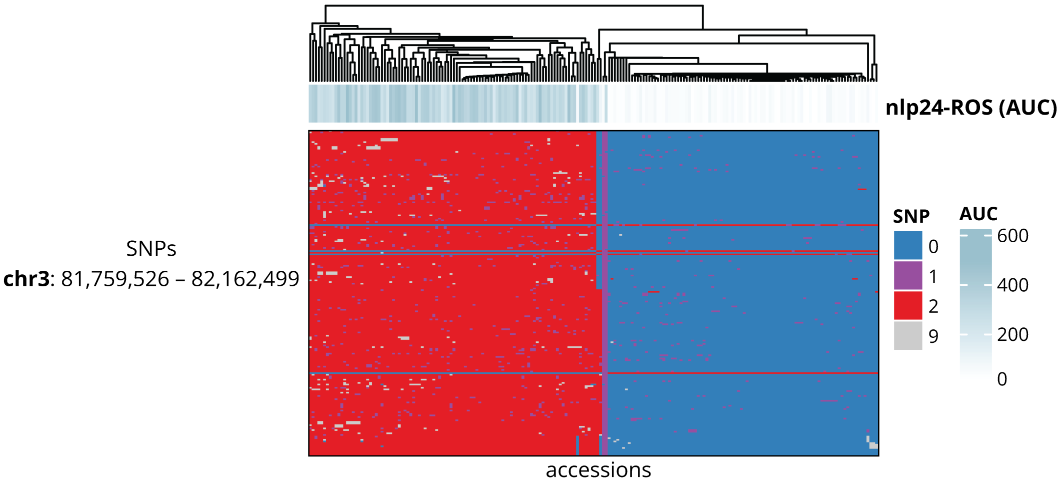


**Supplementary Figure 9**. Single nucleotide polymorphisms (SNPs) across 198 *L. sativa* accessions in the region of the cv. Salinas genome corresponding to coordinates chr3: 81,759,526–82,162,499 (GCF_002870075.4, v11). Rows represent individual SNPs, and columns represent accessions. ROS production after nlp24 treatment is included above (AUC). SNP states are coded as follows: homozygous reference (0), heterozygous (1), homozygous alternative (2), and undetermined (9).

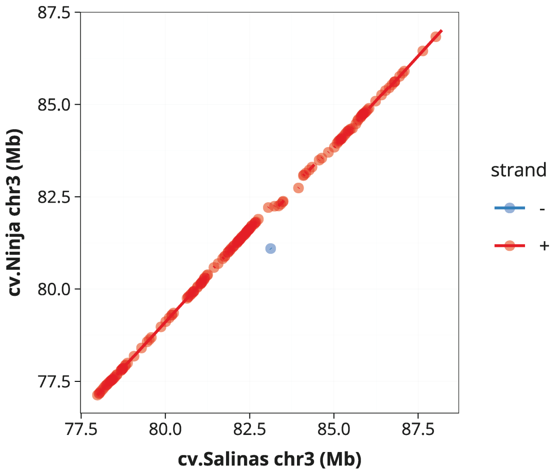


**Supplementary Figure 10**. MUMmer plot of the sequence alignment between the cv. Salinas and cv. Ninja loci shown in figure 3d. Each dot represents a match between the two genomes. Forward and reverse complement matches are shown in red and blue, respectively.

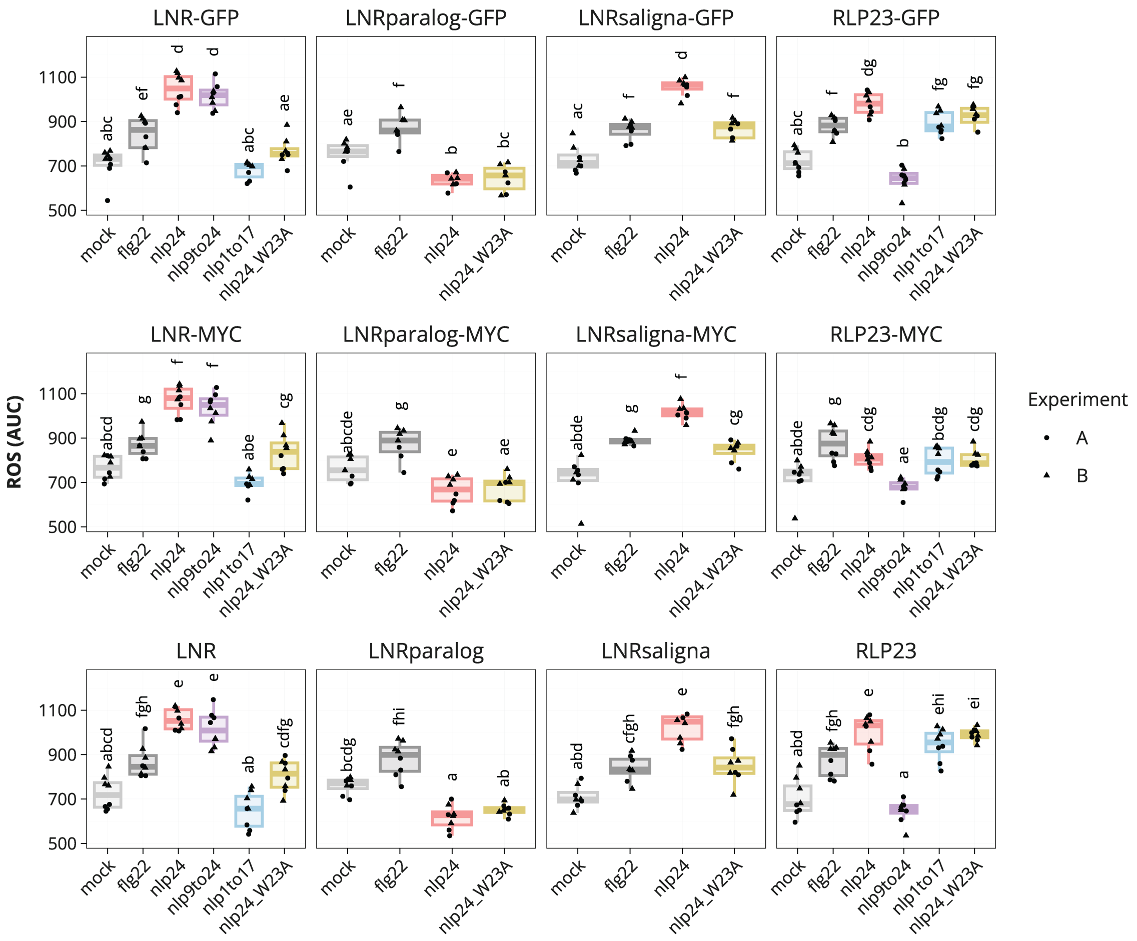


**Supplementary Figure 11.** ROS burst in lettuce cv. Salinas transiently expressing untagged, GFP-, or MYC-tagged versions of *RLP23*, *LNR*, a closely related *RLK* in cv. Salinas (*LNRparalog*), or an ortholog from *L. saligna* (*LNRsaligna*) exposed to different elicitors. Letters represent levels of statistical significance between mean AUC values for all constructs with the same tag (ANOVA, Tukey’s HSD, $p<0.05$; n=7-8; two independent experiments).

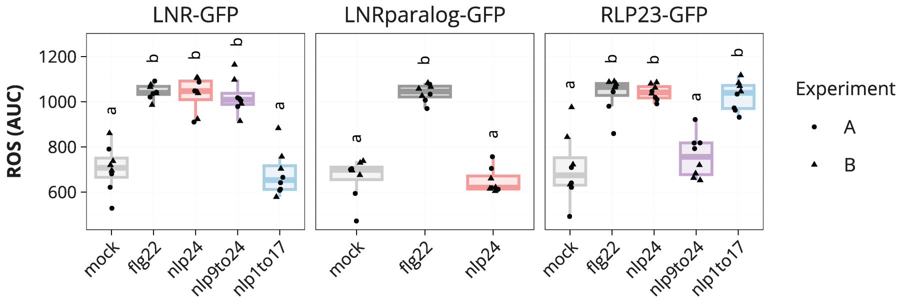


**Supplementary Figure 12.** ROS burst in *N. benthamiana* transiently expressing, GFP-tagged versions of *RLP23*, *LNR*, or a closely related *RLK* (*LNRparalog*), and exposed to different elicitors. Letters represent levels of statistical significance between mean AUC values for all constructs (ANOVA, Tukey's HSD, $p<0.05$; n=7-8; two independent experiments).

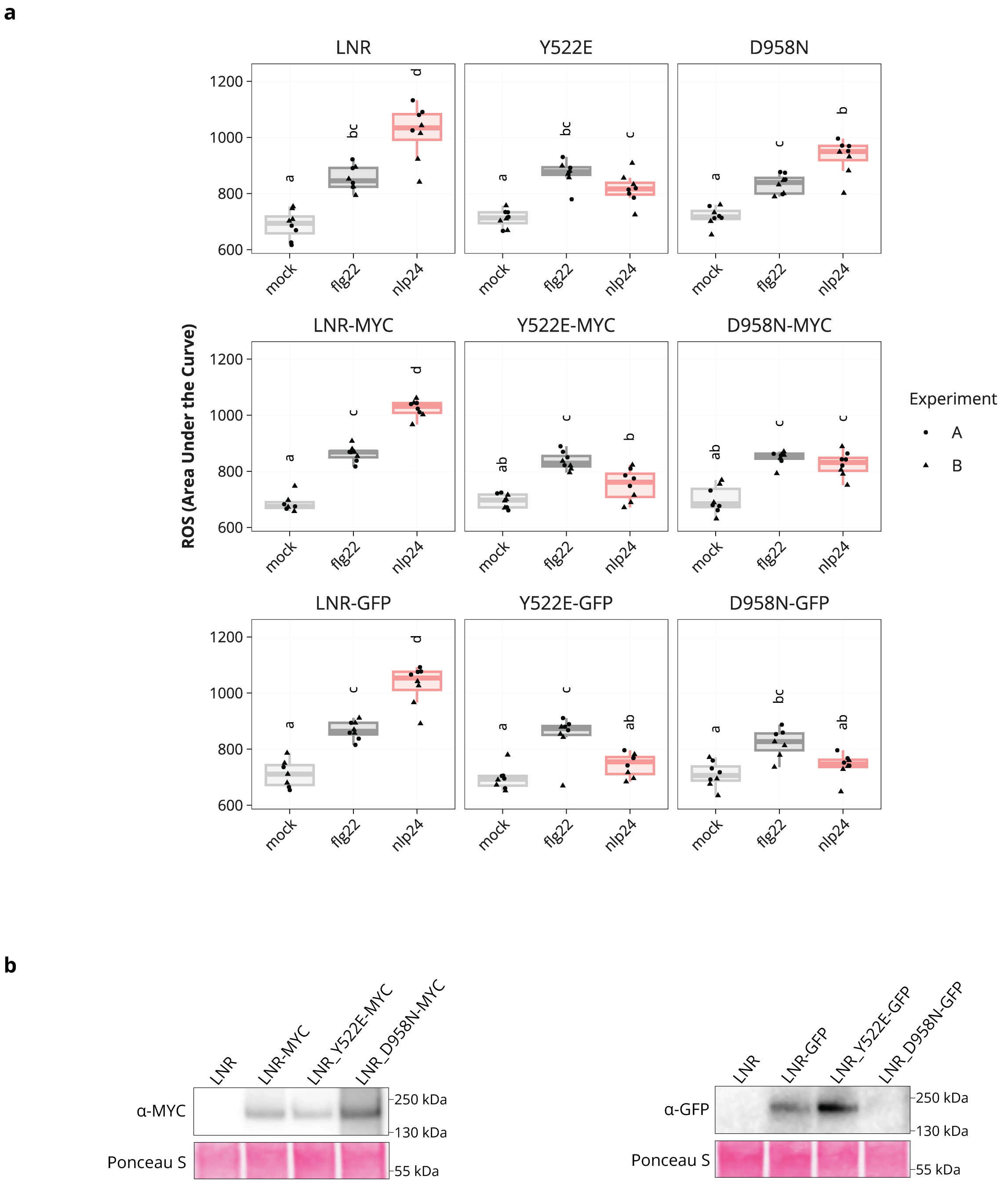


**Supplementary Figure 13. a.** ROS burst in lettuce cv. Salinas transiently expressing untagged, GFP-, or MYC-tagged versions of *LNR*, *LNR*$^{Y522E}$, or *LNR*$^{D958N}$, and exposed to different elicitors. Letters represent levels of statistical significance between mean AUC values for constructs with the same tag (ANOVA, Tukey's HSD, $p<0.05$; n=7-8; 2 independent experiments). **b.** Western blot showing the accumulation of MYC- and mEGFP-tagged LNR and single residue mutants in transient expression assays in lettuce cv. Salinas. Untagged LNR is included as a negative control. Detection was performed with α-MYC and α-GFP antibodies. Ponceau S staining of the blots serves as a loading control. Accumulation was confirmed in an independent experiment.

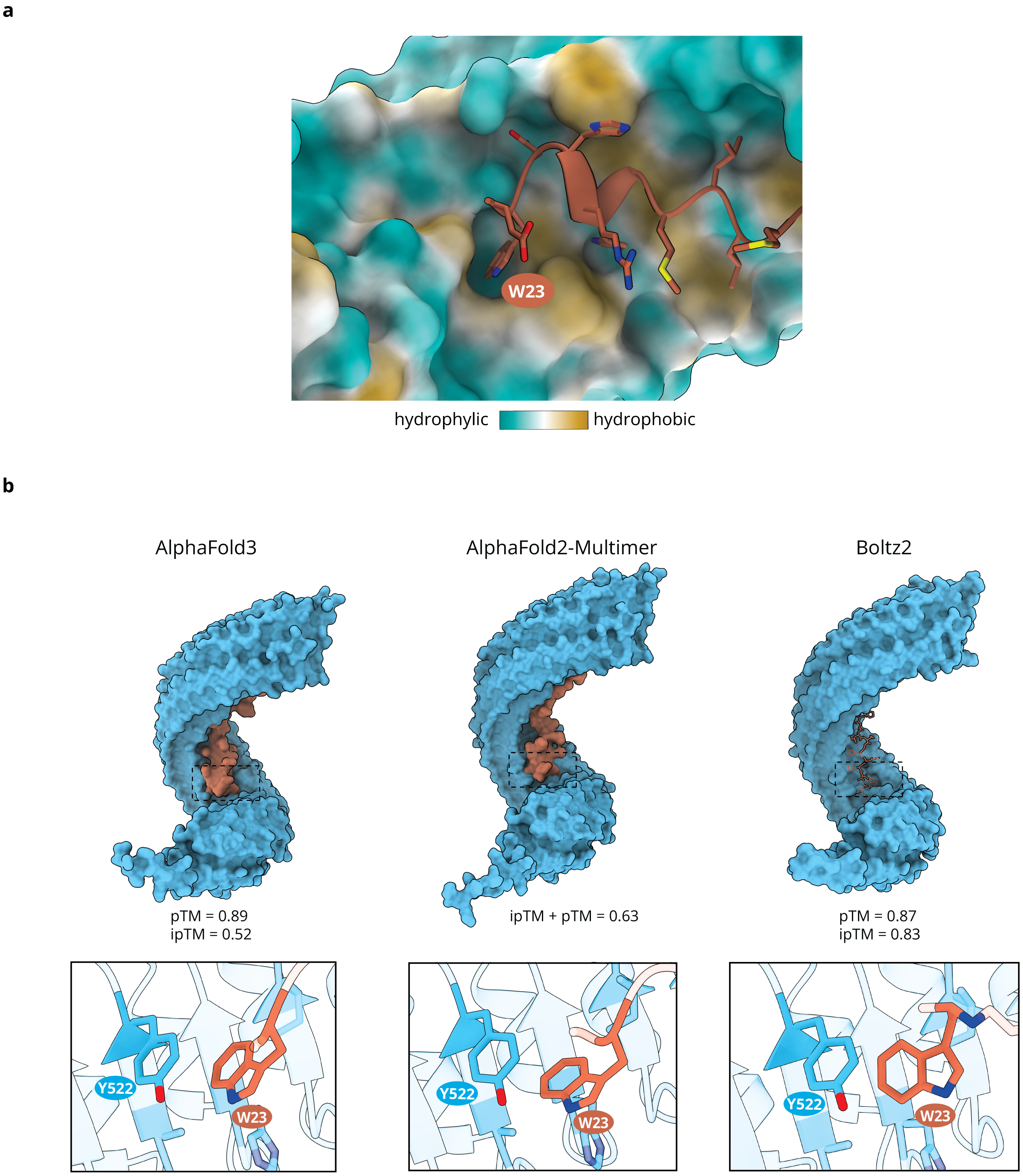


**Supplementary Figure 14. a.** Predicted interaction between nlp24$^{W23}$ and a hydrophobic pocket in the inner concave surface of the LNR ectodomain (AlphaFold3). **b.** LNR-nlp24 predicted interaction by AF3, AF2-Multimer, and Boltz2. nlp24 W23 residue is predicted to insert into a pocket in the LNR surface and closely interact with Y522 by all modeling approaches. pTM, ipTM, and ipTM+pTM values are provided.

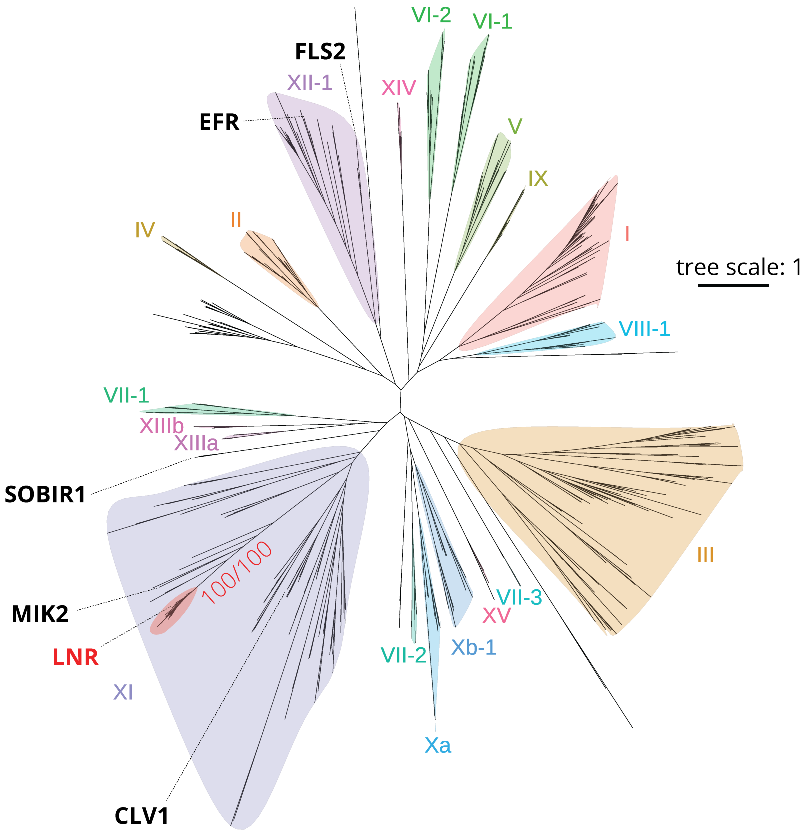


**Supplementary Figure 15**. Phylogeny of LRR-RLKs in *Arabidopsis* and *Lactuca sativa* based on the kinase domain alignment. Proteins mentioned in the main text are highlighted, including LNR and its monophyletic group. Ultrafast bootstrap and SH-aLRT support values are included for the LNR clade (1000 replicates).

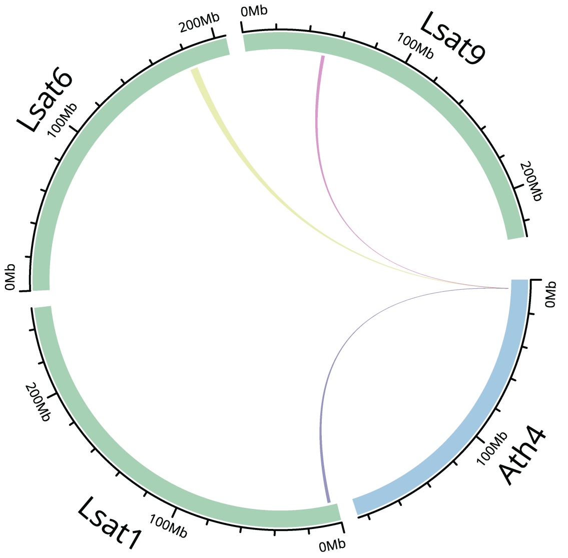


***Lactuca sativa*** **cv. Salinas**
regions with synteny to the *Arabidopsis MIK2* locus

chr1: 5,670,983 - 3,776,992 (reverse)
chr6: 179,437,840 - 184,233,869 (forward)
chr9: 49,351,218 - 51,425,548 (forward)

**Supplementary Figure 16**. Syntenic relations between the *Arabidopsis MIK2* locus (chr4: 5.136.479-6.140.952) and the lettuce cv. Salinas reference genome (GCF_002870075.4). Three syntenic blocks were found on chromosomes 1, 6, and 9. Only the locus on chromosome 9 conserves a *MIK2* copy (*LOC111921259*).

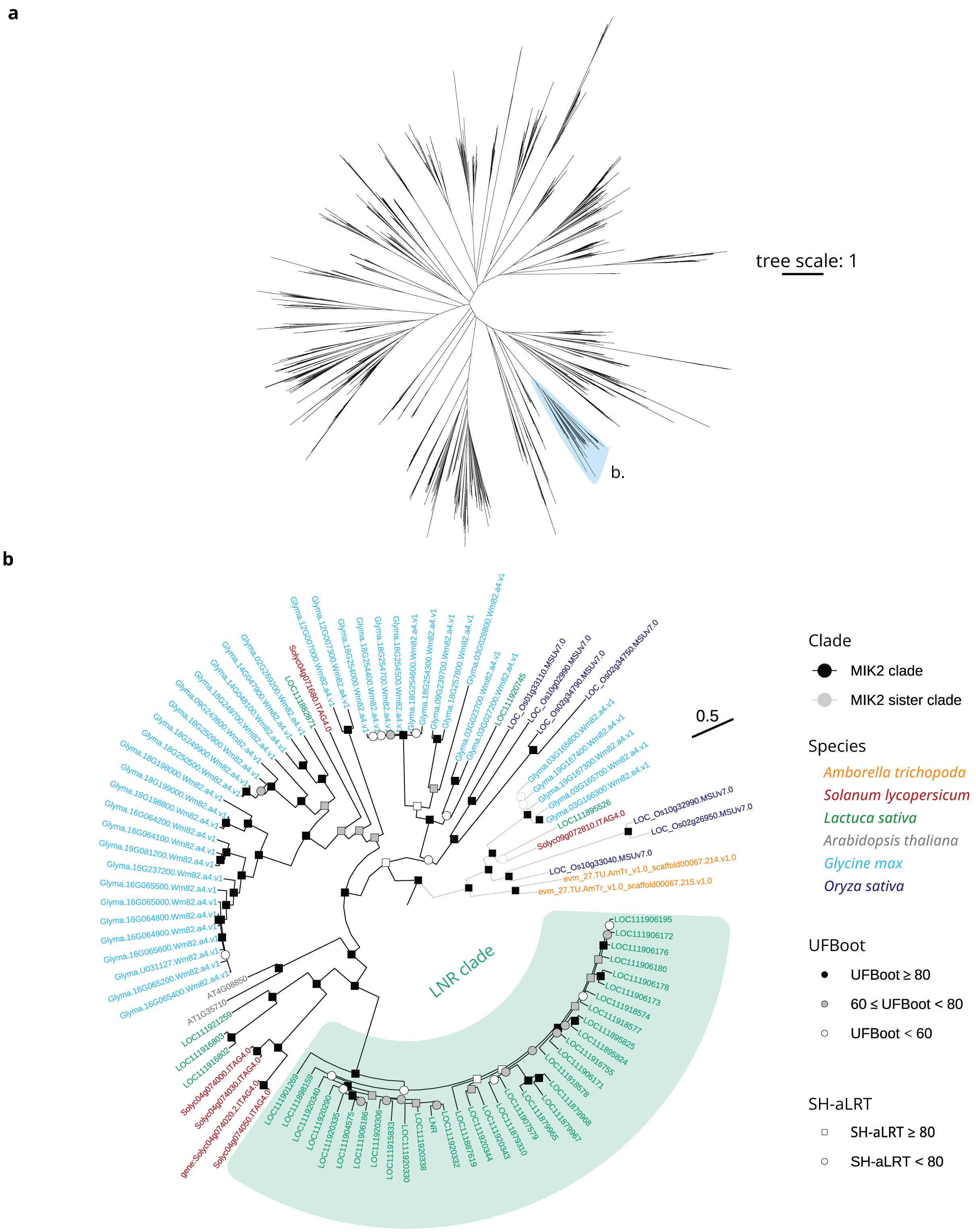


**Supplementary Figure 17**. **a.** Unrooted phylogeny of LRR-RLKs across representatives of major angiosperm lineages. Species included are *Amborella trichopoda*, *Solanum lycopersicum*, *Lactuca sativa*, *Arabidopsis thaliana*, *Glycine max*, and *Oryza sativa*. Branch leading to the MIK2 and the MIK2 sister clade is highlighted. **b.** Expanded view of the MIK2 clade and its sister group. LNR clade within MIK2 clade is highlighted. Node symbols indicate support values, with fill corresponding to ultrafast bootstrap (UFBoot) and shape to SH-aLRT.

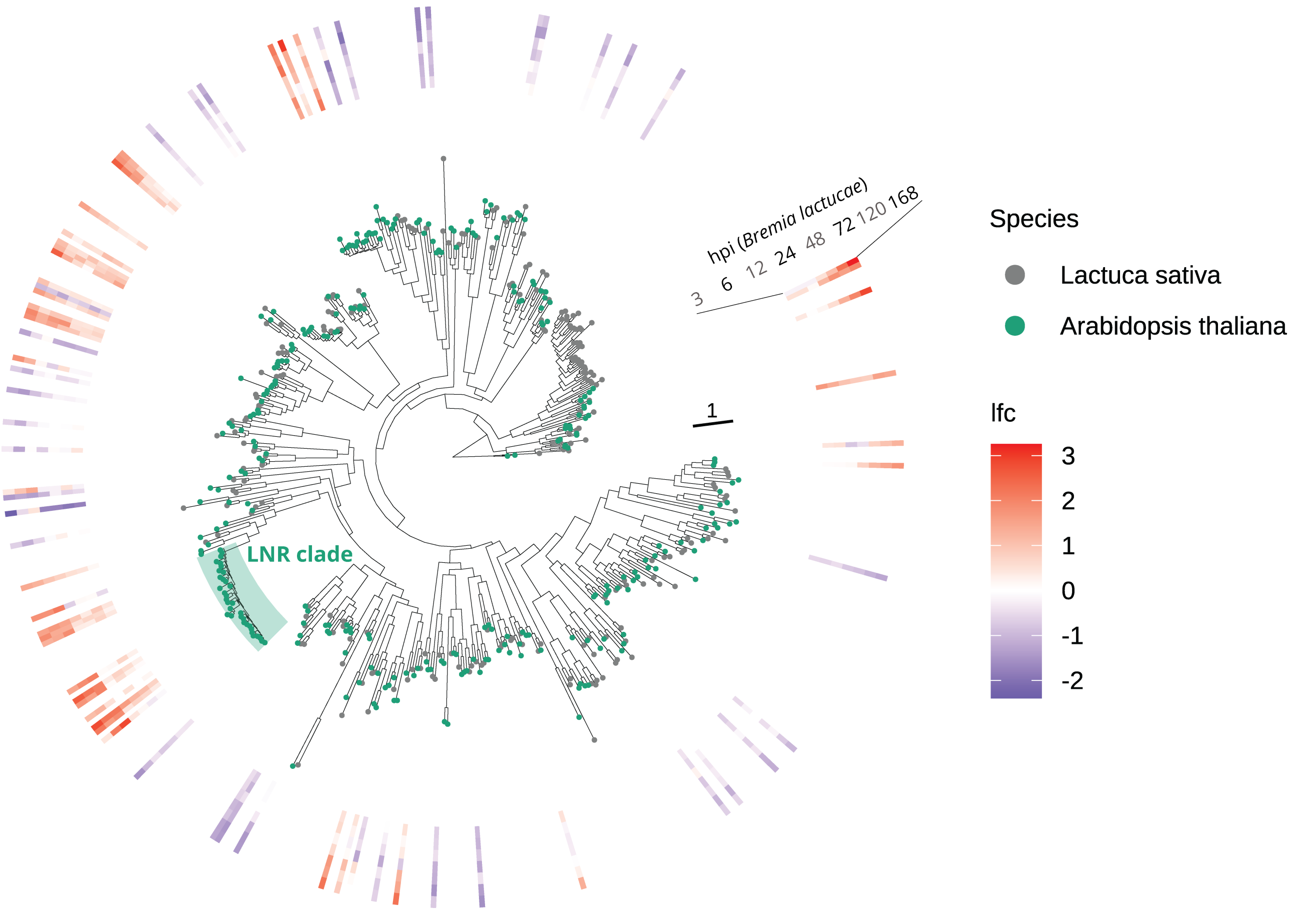


**Supplementary Figure 18.** Phylogeny of LRR-RLKs in *Arabidopsis* and *Lactuca sativa*. Tips are colored based on species. The heatmap shows expression changes ($log_2$ fold change) over time (3h up to 7 days) after *Bremia lactucae* infection (31).

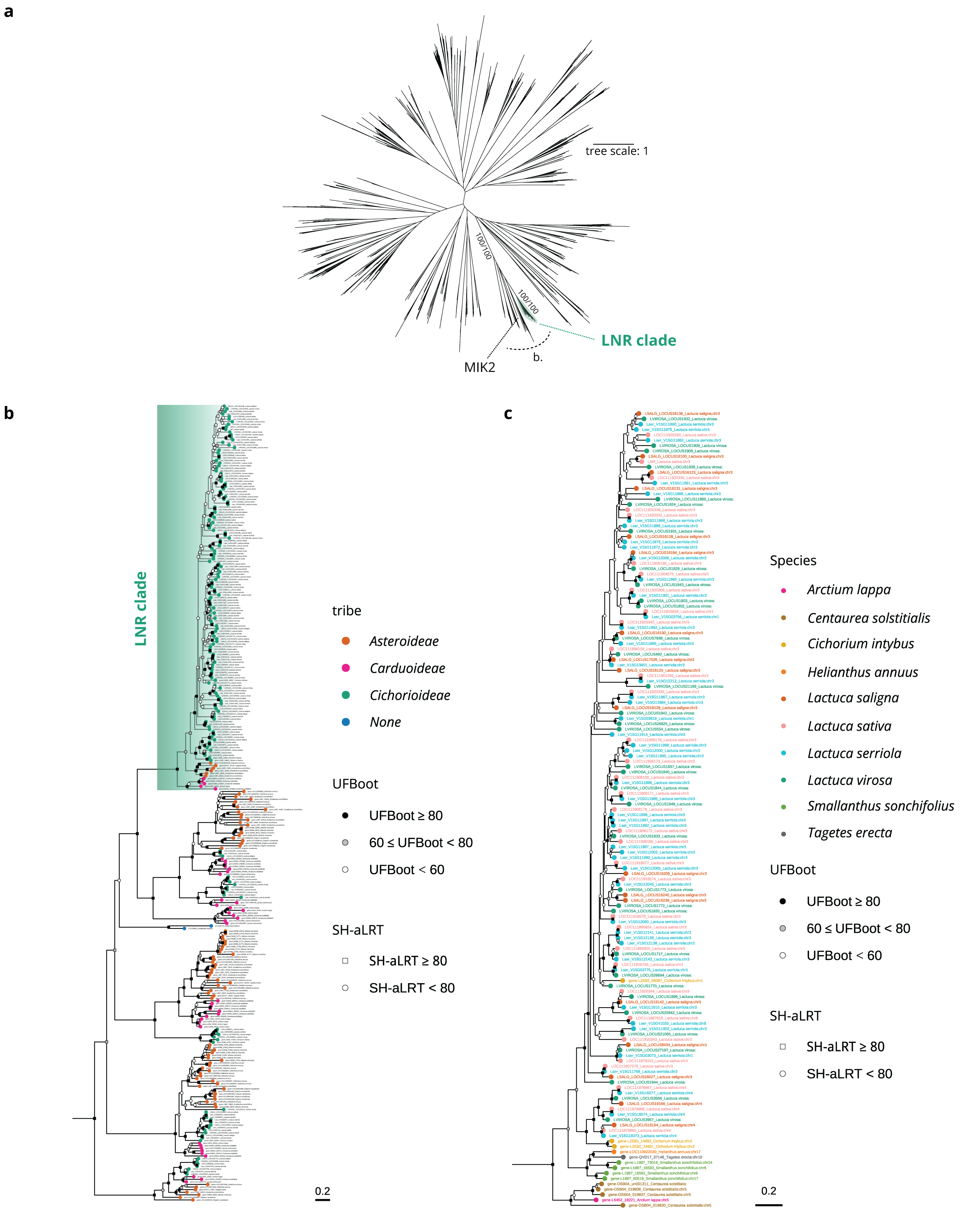


**Supplementary Figure 19**. **a.** Unrooted phylogenetic tree of LRR-RLKs in 13 *Asteraceae* genomes and *Arabidopsis thaliana*. Species that are included are *Lactuca sativa*, *Lactuca serriola*, *Lactuca saligna*, *Lactuca virosa*, *Centaurea solstitialis*, *Cynara cardunculus*, *Arctium lappa*, *Cichorium intybus*,

*Erigeron canadensis*, *Smallanthus sonchifolius*, *Tagetes erecta*, *Mikania micrantha*, *Helianthus annuus,* and *Arabidopsis thaliana*. The maximum-likelihood tree was inferred with IQ-TREE3 (135), and branch supports were assessed with 1000 replicates each of ultrafast bootstrap (UFBoot) and SH-like approximate likelihood ratio test (SH-aLRT). Support values for the LNR clade and the branch leading to it are shown (SH-aLRT/UFBoot). **b.** Zoom-in of the branch that includes MIK2 genes and the LNR clade, colored by tribe. **c.** Close-up view of the LNR clade, colored by species.

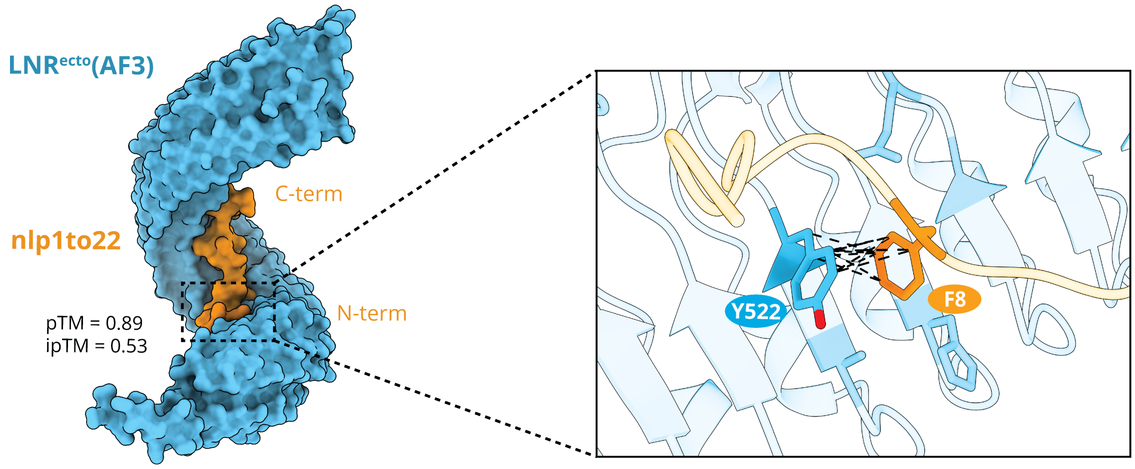


**Supplementary Figure 20.** Interaction between LNR ectodomain and nlp1to22 modeled by AlphaFold3. pTM and ipTM scores are included. Zoom-in of the interaction between LNR$^{Y522}$ and nlp24$^{F8}$ is shown.
